# Immunodominant surface epitopes power immune evasion in the African trypanosome

**DOI:** 10.1101/2021.07.20.453071

**Authors:** Anastasia Gkeka, Francisco Aresta-Branco, Gianna Triller, Evi P. Vlachou, Mirjana Lilic, Paul Dominic B. Olinares, Kathryn Perez, Brian T. Chait, Renata Blatnik, Thomas Ruppert, C. Erec Stebbins, F. Nina Papavasiliou

## Abstract

The African trypanosome survives the immune response of its mammalian host by antigenic variation of its major surface antigen (the Variable Surface Glycoprotein, or VSG). Here we describe the antibody repertoires elicited by different VSGs. We show that the repertoires are highly restricted, and are directed predominantly to distinct epitopes on the surface of the VSGs. They are also highly discriminatory: minor alterations within these exposed epitopes confer antigenically-distinct properties to these VSGs and elicit different repertoires. We propose that the patterned and repetitive nature of the VSG coat focuses host immunity to a restricted set of immunodominant epitopes per VSG, eliciting a highly stereotyped response, minimizing cross reactivity between different VSGs and facilitating prolonged immune evasion through epitope variation.

## Introduction

The African trypanosome (*Trypanosoma brucei spp*) is a strictly extracellular parasite, infection with which elicits a robust antibody response against its Variant Surface Glycoprotein (VSG) coat (Stijlemans et al., 2017; Verdi and Zipkin et al., 2020). This response mediates parasite clearance, but is vulnerable to immune escape from variants that have switched surface coat composition (Mugnier et al., 2015). The VSG protein consists of a large N-terminal domain (NTD, approximately 350 amino acids) and a shorter C-terminal domain (CTD, approximately 100 amino acids), the latter being linked to the membrane via a GPI anchor (Grünfelder et al., 2002). It is speculated that the majority of immune epitopes are located on the tope lobe of the NTD, which is highly accessible to the immune system, however there are no antibody-VSG co-crystal structures available to support this hypothesis (Aresta-Branco and Erben et al., 2019; Bangs, 2018; Zoll et al., 2018), and recent studies have challenged this model (Hempelmann et al., 2021).

The dynamic interplay between VSG-coat-switching by trypanosomes and the B cell response of the host that they evade is central to the maintenance of chronic infection (Silva-Barrios et al., 2018). Nonetheless, very little is known about the nature of the antibody response against the VSG coat. A small number of papers dating back to the 1990s revealed that acute infection with *T. brucei* leads to extraordinarily high levels of IgM in humans and in mice (Verdi and Zipkin et al., 2020). The antibody response has often been characterized as polyclonal and many of those antibodies are autoreactive (Diffley, 1983; Müller et al., 1996; Verdi and Zipkin et al., 2020). However, these data are conceptualized in the context of a prolonged infection that itself is polyclonal in VSG-archetype. An understanding of the response to individual VSG variants in a controlled setting is therefore lacking, since the culmination of multiple separate responses to different VSGs would produce a “polyclonal” B cell output in the greater infection context. Importantly, immunization or exposure to specific VSG surface coats prevents later infection by cells expressing that same coat, indicating that coat-specific antibodies are produced during infection (Campbell et al., 1977; Mugnier et al., 2015).

In order to explore the parameters of the antibody response to *T. brucei*, we examined the antibody repertoires elicited by two specific VSGs. These were chosen to broadly represent the two primary VSG “classes”: VSG2 (a Class-A VSG) and VSG3 (a Class-B VSG) (Carrington et al., 1991). Herein, we demonstrate that the repertoires show restricted V-gene usage which is highly reproducible between animals and experiments. Moreover, we show that small alterations (e.g., point mutations) within VSG surfaces result in distinct repertoires, suggesting that antibodies elicited by VSGs are selective toward specific epitopes on the surface of each VSG, and can discriminate even between single amino acid changes. We speculate that this intense immune-focusing of antibody responses to a very small number of epitopes per VSG may have evolved to reduce cross-reactivity between VSGs and to facilitate prolonged immune evasion.

## Results

### Trypanosome infections lead to rapid plasma cell expansion

To generate the antibody repertoires, we infected animals with each specific VSG-expressing parasite, and either “cleared” the parasites with the anti-protozoan compound diminazene four days after infection (as infection is often rapidly lethal) (Pinger et al., 2017) or allowed the animals to clear the trypanosomes naturally (whenever feasible) after the first peak of parasitemia (at which point the parasites that have switched VSG coats remain and seed the next peak of parasitemia). Initial analyses of infected C57BL/6 mice showed a robust expansion of plasma cells (Figure 1A, Figure S1A). This expansion was common to all infections regardless of whether parasites were cleared by the animals (usually by day 6 or day 7 post infection) or whether they were pharmacologically cleared earlier with diminazene (days 4 and 5), allowing for the maturation of the antibody response until collection (which was uniformly at day 8 post infection – Figure 1B). This is also consistent with known parameters of disease progression (Black et al., 1985; Jayaraman et al., 2019; Mugnier et al., 2015).

**Figure 1:**
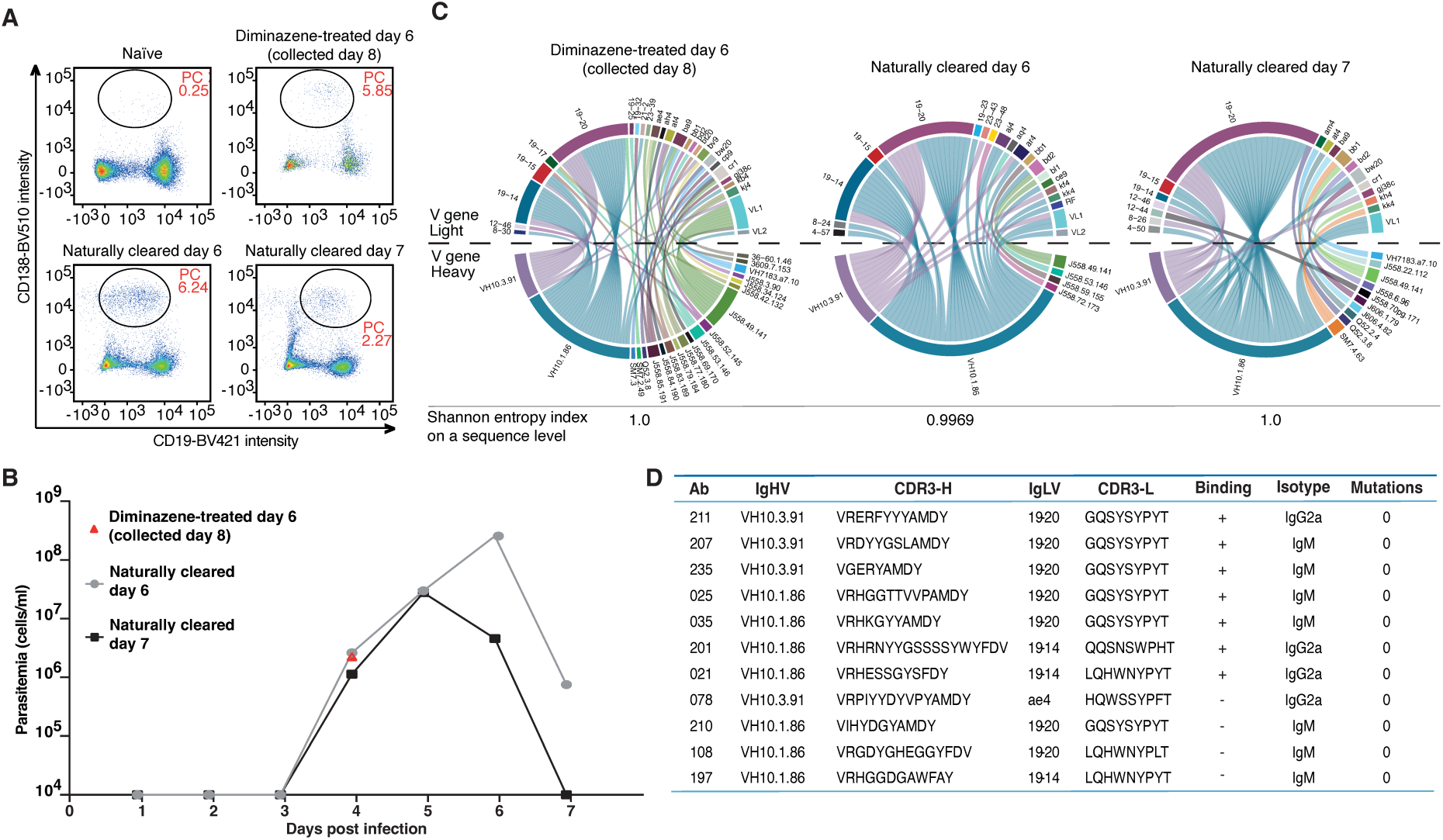
VSG2_WT_ parasites elicit a highly restricted antibody repertoire, defined by a small number of heavy and light chain pairs. **(A)** Percentage of CD138+ plasma cells (PC) within the live lymphocyte population in a non-infected naïve mouse and in infected mice that were treated with diminazene or that naturally cleared the first peak of parasitemia. Plasma cells from this gate were sorted for antibody repertoire analysis. Representative gatings to determine live lymphocytes are shown in Figure S1A. **(B)** Parasitemia infection curves from the 2 mice that naturally cleared the first peak of parasitemia (panel A) at day 6 (black) and day 7 (grey). Both mice were sacrificed at day 7 post-infection. A representative value of parasitemia at day 4 for mice treated with diminazene is indicated in red. **(C)** Circos plots generated for the VSG2_WT_ V gene signatures of two independent mice whose infections were treated with diminazene (left - n=89 pairs) and of two additional mice that naturally cleared the first peak of parasitemia at days 6 (center - n=53 pairs) and 7 (right - n=50 pairs) post infection. Different colors depict each heavy chain variable gene (bottom half of the plot), and each light chain variable gene (top half of the plot). The heavy and light chain variable gene pairings that form the antibodies are illustrated as connector lines starting from the heavy chain genes, providing also information about the frequency of the pairings. Genes from both chains that appeared only once and resulted in single heavy-light pairings, were considered background and were removed from the plot. Shannon entropy shows clonal diversity on a sequence level with a value of 1.0 representing 100% clonal diversity (no clones), while a value of 0.0 corresponding to 0% clonal diversity (only clones). **(D)** Table with representative VSG2_WT_ antibodies elicited in the context of diminazene-treated infections, which are mostly composed of V*H*10 heavy chains paired with 19-14 and 19-20 Vκ light chains. The CDR3 from each antibody is described, indicating that the repertoire is not monoclonal. The original antibody isotype, as well as somatic hypermutations, if any, are also shown.

Typically, the characterization of an antigen-specific antibody repertoire benefits from the “baiting” of antigen specific B cells using the antigen itself conjugated to a fluorophore (PE or APC) (Scheid et al., 2009; Tiller et al., 2008). However, plasma cells are notoriously hard to bait as they downregulate surface expression of Ig and rather secrete the vast majority of the antibody they produce, so that very little antibody remains surface-bound and accessible. We therefore proceeded to sort all plasma cells according to surface marker expression (CD138+/CD19^lo^) (Pracht et al., 2017) into 384-well plates. The antibody repertoire of plasma cells was then assembled after two rounds of single cell PCR and paired heavy (IgH) and light (Igκ/λ) gene sequence analysis (Tiller et al., 2008).

### The VSG2_WT_ repertoire is characterized by a small number of heavy and light chain variable region genes

We first focused on the repertoire elicited by VSG2-coated parasites. Surprisingly for a non-baited repertoire, plasma cells elicited from mice infected with VSG2 were dominated by cells carrying only four VH+VL chain pairings (VH10.1 or VH10.3 with VL19-20 or VL19-14, Figure 1C). These combinations, which are absent in naïve mice (Figure S1B), formed the plurality of the repertoire when mice were pharmacologically treated to remove trypanosomes four days after infection, but they became increasingly prevalent in repertoires of animals that cleared trypanosomes naturally a day before collection, and even more dominant in repertoires of animals that cleared the infection the day we collected plasma cells (Figure 1C). This suggests a rapid and robust response to the parasite, stereotyped and highly reproducible between mice, that may become increasingly diversified as more antigens become available after parasite lysis.

We reasoned that the repertoire we collected closest to natural clearance would contain the B cells likely expressing the most potent anti-coat-specific antibodies and proceeded to analyze it further. While dominated by cells expressing two distinct immunoglobulin (Ig) heavy and Ig light chain gene pairs, the B cells elicited by VSG2-coated *T. brucei* were not monoclonal: each VH10 heavy and 19-20 or 19-14 light chain V region was joined to a set of DJ or J gene segments representing a wide swath of CDR3 variants (Figure 1D). This lack of clonal expansion is also evident from the Shannon entropy index, which is a statistical sequence-based measure of clonality (Figure 1C, bottom of each circos plot).

We therefore picked a small number of heavy and light chain pairs from the overrepresented V gene segments identified and proceeded to express them recombinantly as soluble antibodies in 293T cells. Indeed, 7 out of 11 of these antibodies are capable of binding live, VSG2-coated *T. brucei*, but not parasite cells coated with a different VSG protein (Figure 1D, Figure 3C). Antibodies that did not use these V segments were not able to bind VSG2- coated parasites (Figure S2). We conclude that the VSG2-elicited plasma cell repertoire represents an oligoclonal expansion primarily of four variable region defined pairs, which are mostly unmutated (Figure 1D). The V gene signature that defines this repertoire is stereotyped between mice (Figure 1C), suggesting both that the VSG2-specific paratope is likely located within highly-specific variable regions of the antibody protein, and that these antibodies are likely generated against a common immunodominant epitope.

### The structure of VSG2_WT_ contains a calcium binding pocket, which defines the immunodominant epitope

Epitope identification is often achieved by structural analyses of repertoire-defined antibodies together with the antigen that elicited them (Triller et al., 2017). We therefore sought to crystallize VSG2 (specifically, the larger, surface-exposed N-terminal domain of the VSG, or “NTD”) together with VSG2-specific Fab-fragments. All crystals obtained were of VSG2 alone, however (Figure S3C, Table S1), likely because most antibodies elicited by *T. brucei* infection are of the IgM isotype (Figure S4, A and B) (Verdi and Zipkin et al., 2020) and are likely of low affinity (but high avidity). These antibodies are thus unlikely to bind with high-affinity when produced recombinantly as an IgG (or Fab) fragment (as current methods for the recombinant production of pentameric IgM are poorly optimized). Serendipitously, the crystals did allow us to re-solve the structure of VSG2 to much higher resolution compared to previous work (Freymann et al., 1990; Metcalf et al., 1987), with unexpected findings. The protein structure of VSG2 from these new crystals is nearly identical to that from the 1990 crystal form, with a root-mean-square deviation of 0.87Å over 1420 atoms of a monomer in the dimer (Figure 2A, Figure S3A). However, in later stages of model refinement, unusually large electron density around solvent molecules was modeled effectively with calcium (Figure 2, B and C), an ion not present in older structures likely due to the use of chelating agents in the purification from animal blood (Cross, 1975, 1984). Calcium is octahedrally coordinated in VSG2 through a network of oxygen bonds: four from three consecutive amino acid side-chains at the end of a β-strand (bidentate-D208, N209, and D210, henceforth the “DND motif”), the carbonyl-oxygen from G155 in an opposing loop, and three water molecules in the solvent shell above the protein (each water coordinated by other side-chain and main-chain contacts from the protein, Figure 2C).

**Figure 2:**
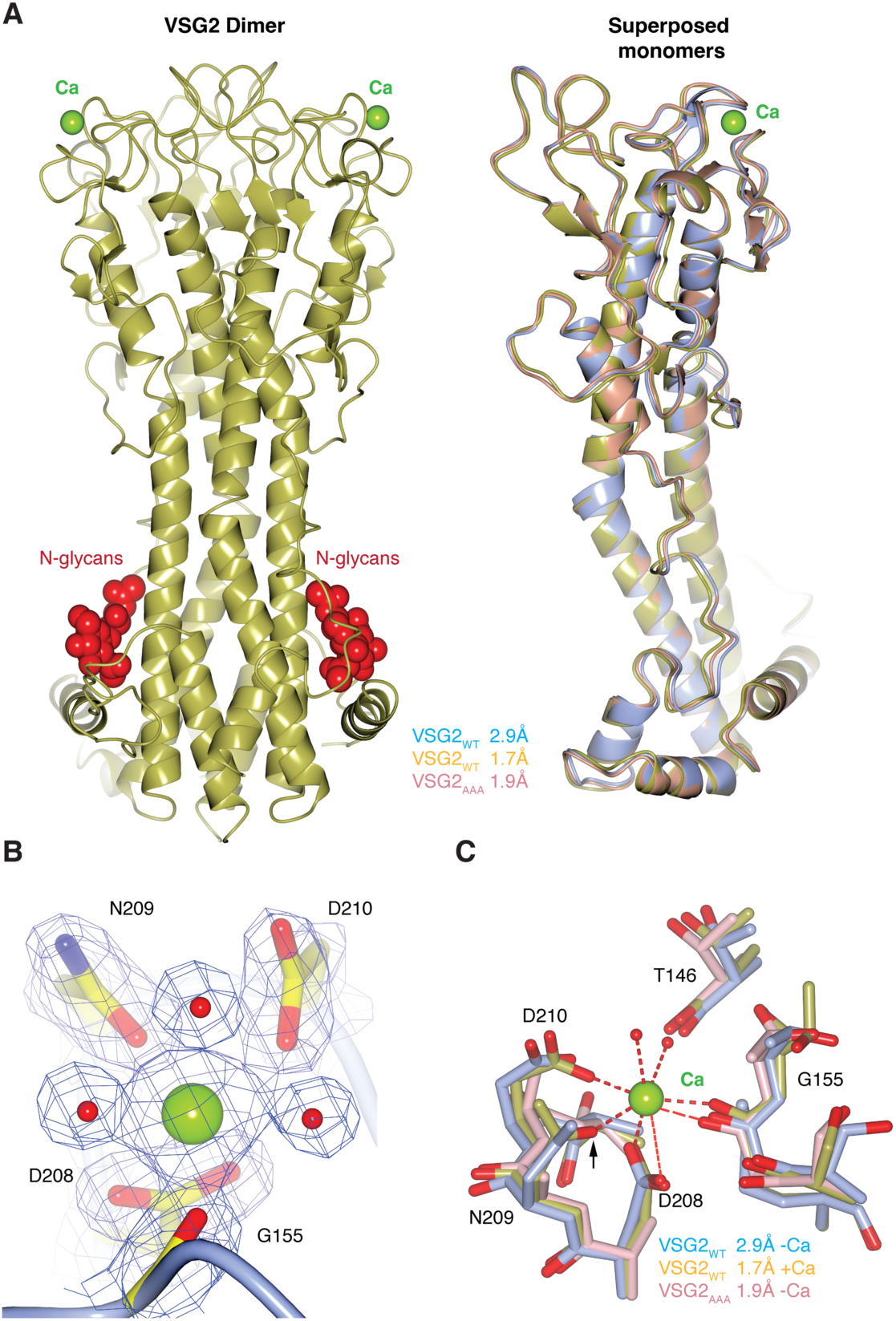
The resolved crystal structure of VSG2_WT_ reveals the existence of a surface-exposed calcium binding pocket. **(A)** Left - Crystal structure of VSG2_WT_. The homodimer is shown as a ribbon diagram colored in gold, N-linked glycans and calcium atoms are displayed as red and green space-filling atoms, respectively. Right – Structural alignment of monomers of VSG2_WT_ (from left panel, in gold), VSG2_WT_ (from (Freymann et al., 1990), in blue) and VSG2_AAA_ (in pink), highlighting the coordination of the calcium atom. **(B)** Close-up of the coordination of calcium binding pocket with corresponding electron density maps. Water molecules and calcium atoms are displayed as red and green space-filling atoms, respectively. **(C)** Superposition of the calcium binding pockets of the 3 structures presented in (A). Water molecules and calcium atoms are displayed as red and green space-filling atoms, respectively.

There is little change in the fold or amino acid side-chain positioning between the calcium bound and free structures of VSG2 (Figure 2, A-C), indicating that the pocket does not undergo any significant conformational changes upon calcium binding. The presence of calcium was verified both by mass spectrometry as well as isothermal titration calorimetry (Figure S5, A-D). The calcium binding pocket in VSG2 appears to be unique, not closely resembling any published structures in the PDB (e.g., DALI homology search) (Holm, 2020). The pocket occurs near the “top” of the NTD of VSG2, slightly sheltered from the uppermost surface (Figure 2A). A triple mutation of the DND motif to alanine (VSG2_AAA_) abrogates calcium binding (Figure S5, E-G) without any significant conformational changes (Figure 2, A and C, Figure S3, A and D), although electrostatic surfaces generated from the three structures show an expected change to a more neutral charge distribution in the pocket region of the mutant (Figure 3A).

**Figure 3:**
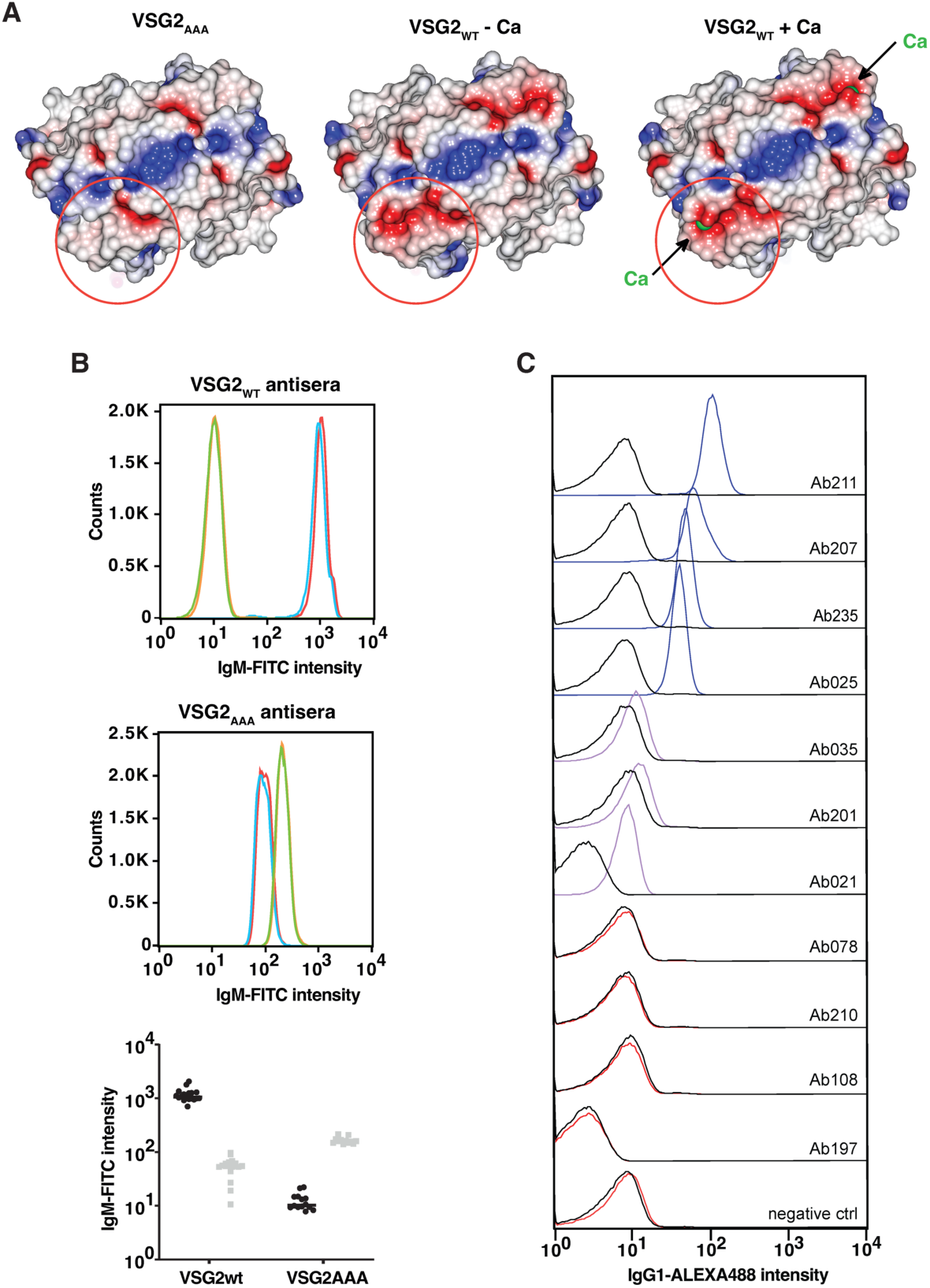
Residues surrounding the calcium binding pocket of VSG2_WT_ define an immunodominant epitope. **(A)** Molecular surfaces of VSG2_WT_ + calcium (this study), VSG2_WT_ - calcium (from (Freymann et al., 1990)) and VSG2_AAA_. The VSGs are oriented looking ‘down’ on the top lobe of the protein, the orientation rotated 90° about a horizontal axis in comparison to upper panels. The surfaces are colored by relative electrostatic potential (blue is basic/positively charged, red is acidic/negatively charged, and white is neutral). **(B)** Histograms reflecting binding intensities of antisera collected from mice infected with VSG2_WT_ and VSG2_AAA_ parasites (top two panels, respectively) with VSG2_WT_ and VSG2_AAA_ parasites (blue and red correspond to 2 independent clones of VSG2_WT_ parasites; green and orange correspond to 2 independent clones of VSG2_AAA_ parasites). Bottom panel shows quantification of binding intensities of antisera collected from mice infected with VSG2_WT_ (black dots) and VSG2_AAA_ (grey dots) parasites with VSG2_WT_ and VSG2_AAA_ parasites (median from n=12-18). **(C)** Histograms reflecting binding intensities of the antibodies presented in Figure 1D with wild type parasites (colors) and AAA-mutant parasites (black). Different colors are meant to distinguish differential binding intensity: red – no binding; purple – poor binding; blue – good binding. The gating strategy is shown in Figure S1C.

Unexpectedly, despite the high similarity between the wild type and mutant structures, antisera elicited by infection with VSG2_WT_ coated trypanosomes do not bind trypanosomes expressing the minor variant represented by the VSG2_AAA_ mutant (Figure 3B). Furthermore, our recombinantly expressed antibodies specific to the VSG2_WT_ coat were unable to bind the VSG2_AAA_ mutant coat (Figure 3C, Figure S2), despite the fact that these mutations did not cause conformational changes in VSG2 (Figure 2, A and C, Figure S3A). This suggests that the VSG2 epitope recognized by both antisera and monoclonal antibodies is located within the region defined by the DND motif and conversely, that antibodies to VSG2-coated trypanosomes are capable of a high degree of epitope discrimination.

### The VSG3_S317A_ repertoire is defined by a signature light chain V gene (gn33), absent from the wild type

To assess whether this combination of epitope immunodominance and antibody discrimination characterizes the epitope space of other VSGs, we analyzed antibody repertoires from plasma cells collected from animals infected with VSG3-coated trypanosomes. VSG3 is a member of a different VSG class with significant structural and post-translational modifications distinguishing it from VSG2 (Class A vs Class B VSGs, respectively) (Carrington et al., 1991). Previously, we had solved the crystal structure of the VSG3 NTD, leading to the unexpected identification of an *O*-linked glycosylation of a serine residue (S317, heterogeneously modified with 0-3 hexose residues) located in the center of the top surface of the protein (Pinger, Nešić and Ali, et al., 2018). Mutation of that serine to an alanine residue ablated glycosylation, reduced parasite virulence, and enhanced the effectiveness of the host immune response (Pinger, Nešić and Ali, et al., 2018). At the time we hypothesized that this was due to the tendency of *O*-linked sugars to adopt multiple conformational states (Lisowska, 2002), thus further diversifying the epitope space of the VSG3 coat. Therefore, to assess whether VSG3 coated trypanosomes lacking this post-translational modification could also elicit a stereotyped immune response toward an immunodominant epitope (such as we have seen for VSG2-coated parasites), we first infected mice with S317A mutant parasites (VSG3_S317A_), and analyzed the resulting plasma cell antibody responses either after natural clearance or after treatment with diminazene. As was the case for VSG2 coated parasites, VSG3_S317A_ coated parasites were also capable of reproducibly eliciting a restricted B cell repertoire in multiple distinct infections (Figure 4A). However, this repertoire was now defined by a major light chain V gene (VL-gn33) pairing with a larger number of heavy chains (Figure 4A). Again, recombinant expression of a number of these heavy and light chain pairs as secreted IgG1s in 293T cells show that many of the antibodies elicited by VSG3_S317A_ coats are capable of binding their cognate parasites (Figure 4, B and C, Figure S6, A and B). Although the common light chain variable gene (gn33) is clearly important for binding, the proper heavy chain is also necessary, since pairing of the gn33 light chain with a different VH can result in a loss of binding (Figure S6, C and D). Whether the common light chain plays a role in increasing antibody binding to the repetitive coat remains to be investigated.

**Figure 4:**
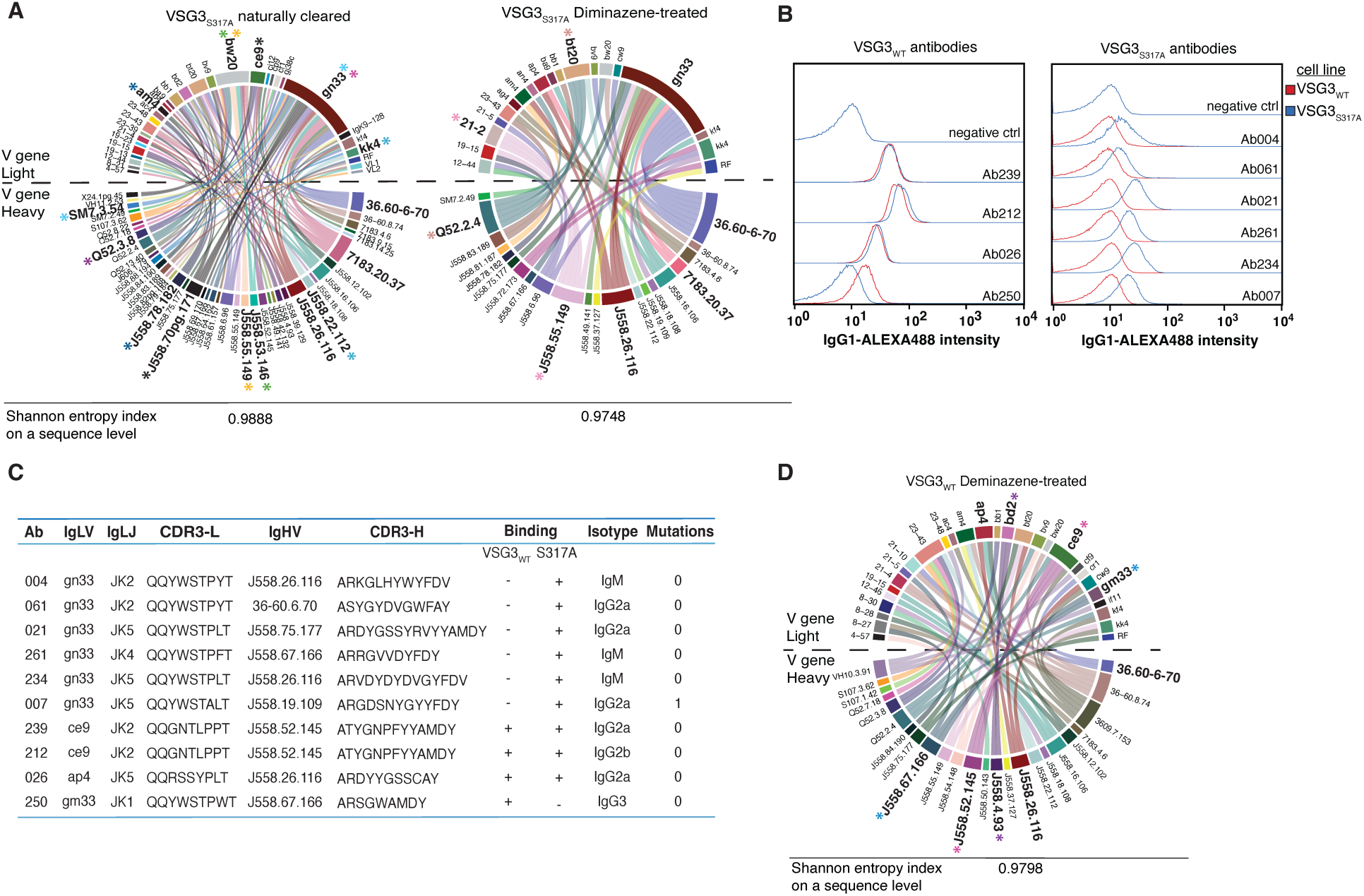
VSG3_S317A_ parasites elicit a restricted B cell response, with preferential utilization of a light chain Vκ gene (gn33). **(A)** Circos diagrams generated for the VSG3_S317A_ V gene signatures of four mice infections, two naturally cleared (left - n=114 pairs) and two diminazene treated (right - n=48 pairs). Different colors depict each heavy chain variable gene (bottom half of the plot), and each light chain variable gene (top half of the plot). The heavy and light chain variable gene pairings that form the antibodies are illustrated as connector lines starting from the heavy chain genes, providing also information about the frequency of the pairings. Genes from both chains that appeared only once and resulted in single heavy-light pairings, were considered background and were removed from the plot. Colored-asterisks depict individual B cells that correspond to clones (i.e., share the same V heavy and V light gene segments but also the same DJ or J genes and the same CDR3 length and sequence). Shannon entropy shows clonal diversity on a sequence level, a number of 1.0 shows 100% clonal diversity (no clones), while a value of 0.0 corresponds to 0% clonal diversity (only clones). **(B)** Left: FACS analysis of heavy and light chain pairs selected from the VSG3_WT_ repertoire, produced recombinantly as IgG1 antibodies. The plots depict antibody binding to VSG3_WT_ (red) or VSG3_S317A_ (blue) *T. brucei* cells. Parasites were first stained with each antibody supernatant, and counter stained with (mouse anti-human) AlexaFluor488-IgG1 secondary antibody. The gating strategy is shown in Figure S1C. Staining with supernatants from untransfected cells (no plasmid) serves as a negative control. All data are normalized to mode. Right: FACS showing the same experimental setup but now for binding of recombinant antibodies selected from the VSG3_S317A_ repertoire. **(C)** Table of all the VSG3_S317A_ antibodies with gn33 as a light chain that bound to the cognate and WT cells, as well as of all the VSG3_WT_ antibodies and their binding to cognate and mutant cells. The (+) symbol indicates binding while the (-) lack of binding. V and J segments as well as the CDR3 of both heavy and light chain genes are shown, along with the binding to the respective parasites. The original antibody isotype, as well as somatic hypermutations, if any, are also shown**. (D)** Circos plots generated for the VSG3_WT_ V gene signatures from two mice whose infections were treated with diminazene (n=53 pairs), as described in (A).

We also analyzed the resulting plasma cell response after infection with VSG3_WT_ followed by treatment with diminazene (as the high virulence of the strain is incompatible with natural clearance). We found that the light-chain restriction seen with VSG3_S317A_ elicited repertoires is now entirely abolished, strongly suggesting that the addition of up to 3 *O*-linked hexoses per VSG3 molecule functions to diversify the presumed immunodominant epitope presented by the VSG3_S317A_ coated parasite (Figure 4D). Furthermore, the addition of sugars entirely prevented the major VL (gn33) carrying antibody pairs from binding VSG3_WT_ coated parasites (Figure 4, B and C), suggesting that antibodies elicited to the parasite coat are focused near-exclusively on the epitope that contains the *O*-linked hexoses. Conversely, of the four antibodies we reconstituted and tested that were elicited by the wildtype (VSG3_WT_) infection, three were able to bind the sugarless coat, and one was selective to the sugar on S317 (capable only of binding the wildtype) (Figure 4, B and C).

### The anti-VSG3 repertoire is highly sensitive to a prominent immunodominant epitope on the surface of VSG3

A comparison of mutant and wild type structures verified that the protein surface between the two VSG3 variants was identical (Figure 5A) but yielded a surprise in the form of an additional sugar in a neighboring serine (S319). We confirmed this in the VSG3_WT_ structure by re-solving it (Figure 5, A and B, Figure S3, B and E, Table S2). S319 is part of the surface-exposed loop harboring S317. In a previous study (Pinger, Nešić and Ali, et al., 2018), both by crystallography and mass spectrometry, glycosylation of S319 was not clearly present (likely due to intrinsic lability of the moiety). However, here we were able to observe strong density for the sugar on S319 in the VSG3_WT_ and VSG3_S317A_ mutant crystal structures, as well as to confirm the presence of hexose groups on S317 and S319 by mass spectrometry (Figure S7). Subsequent determination of the crystal structures of the S319A mutant as well as the double mutant (combining S317A and S319A, or “SSAA”) did not reveal any significant structural differences when compared to VSG3_WT_ or VSG3_S317A_ (Figure 5, A and B, Figure S3, B and F-H).

**Figure 5:**
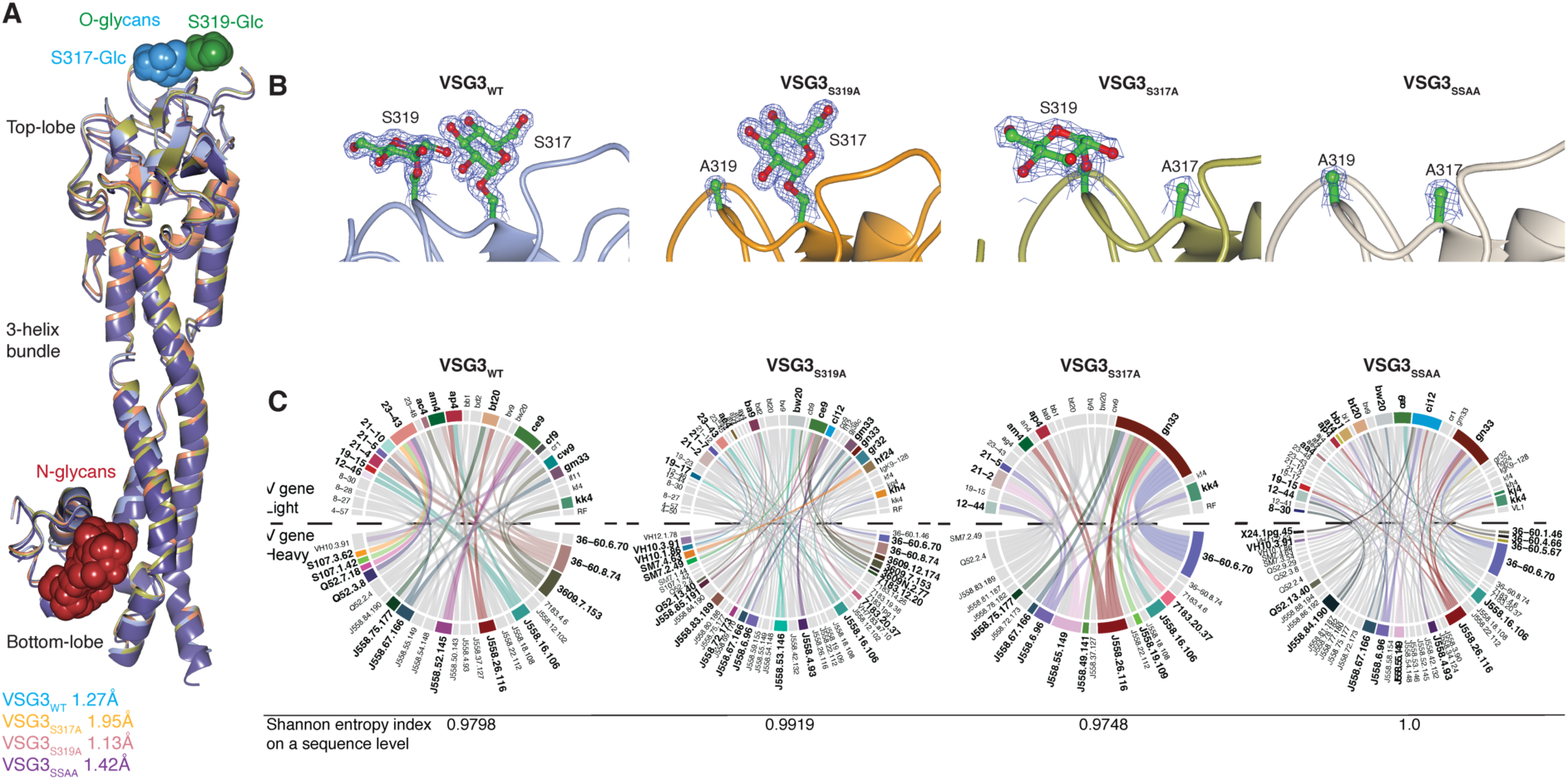
Nearly identical VSG3 variant coats elicit distinct V gene signatures. **(A)** Overlaid crystal structures of all four VSG3 monomers, shown as ribbon diagrams and colored in blue (WT), gold (S317A), salmon (S319A) and purple (SSAA). The N-glycans are represented as red spheres on the bottom lobe, while the *O*-linked sugars as blue (S317-sugar) and green (S319-sugar) spheres on the top lobe. **(B)** Electron density maps maps (2Fo-Fc, contoured at 1σ) focused on the presence or absence of the S317 and S319 *O*-Glucose molecules for all individual proteins as indicated in the labeling. **(C)** Circos plots of the diminazene-treated plasma cell antibody repertoires (V signatures shown only) of WT (n=53 pairs), S317A (n=48 pairs), S319A (n=109 pairs) and SSAA (n=102 pairs). Different colors depict each heavy chain variable gene (bottom half of the plot), and each light chain variable gene (top half of the plot). The heavy and light chain variable gene pairings that form the antibodies are illustrated as connector lines starting from the heavy chain genes, providing also information about the frequency of the pairings. Genes from both chains that appeared only once and resulted in single heavy-light pairings, were considered background and were removed from the plot. The antibody pairs picked for validation are presented in color, while all other genes are in gray. Shannon entropy shows clonal diversity on a sequence level (a value of 1.0 shows 100% clonal diversity (no clones), while a value of 0.0 corresponds to 0% clonal diversity (only clones)).

Following the experimental protocol that we had used to assess the quality of the repertoires elicited by VSG3_WT_ and VSG3_S317A_, we also generated plasma cell antibody repertoires elicited by VSG3_S319A_ and VSG3_SSAA_, (Figure 5C) and tested those for their ability to bind either their cognate parasites or to cross-react with the wild type and other mutants. We found that VSG3_SSAA_ parasites elicited repertoires very similar to those elicited by the VSG3_S317A_ mutant alone (e.g., dominant presence of the gn33 light chain V gene and 30-60.6.70 or J558.26.116 heavy chain gene), whereas VSG3_S319A_ single mutants elicited broader repertoires that were very similar to those elicited by VSG3_WT_ (no gn33 dominance for example) (Figure 5C). In turn, this suggests that the VSG3_WT_ repertoire is strongly influenced by the one-to-three hexoses linked to S317 (Pinger, Nešić and Ali, et al., 2018), but negligibly by the glycans linked to S319; similarly, that the S317A determined repertoire is indifferent to the presence of the sugar on S319, as the repertoire does not change in the double mutant.

We then tested recombinant antibodies that were elicited after infection with each of these variants for the ability to cross-react with each of the other variants, we identified four classes of binders:

a. Antibodies that could bind all four variants (like ab239 or ab222, Figure 6, A and B); these were elicited by both VSG3_WT_ and VSG3_SSAA_ suggesting that they bind a common peptide epitope at a distance from the epitope defining sugar.
b. Antibodies that bound only the epitope-defining sugar-containing variants VSG3_WT_ and VSG3_S319A_ (like ab250, Figure 6, A and B) but not the other two variants; suggesting that they bind the sugar containing epitope but not the same epitope without the carbohydrate (and are therefore potentially sugar-selective);
c. Antibodies that bound only the variants that lacked the epitope-defining sugar at S317 (VSG3_S317A_ and VSG3_SSAA_, like ab021); suggesting that they bind the same peptide epitope defined by the sugar but are inhibited by the presence of the sugar in the other two variants.
d. Finally, antibodies that only bound the VSG3_S317A_ variant (e.g., ab234), potentially suggestive of an additional minor epitope partly defined by the S319 sugar.

**Figure 6:**
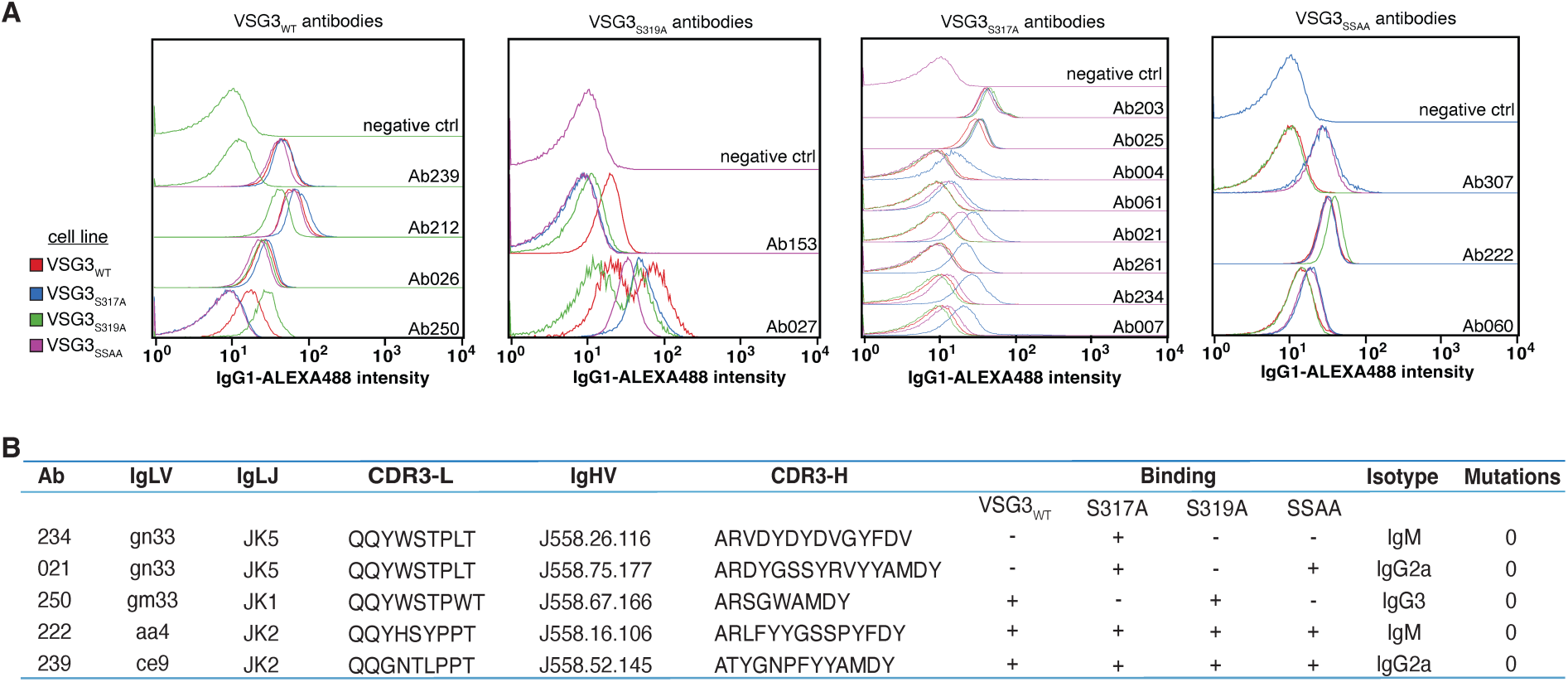
Antibodies elicited by VSG3 variants fall into four classes based on their antigen binding capacity. **(A)** FACS data showing the binding of the validated antibodies, secreted as IgG1s, to VSG3_WT_ (red), VSG3_S317A_ (blue), VSG3_S319A_ (green) and VSG3_SSAA_ (purple) - covered trypanosomes. Parasites were first stained with each antibody supernatant, followed by staining with a (mouse anti-human) AlexaFluor488-IgG1 secondary antibody. The gating strategy is shown in Figure S1C. Staining with supernatants from untransfected cells (no plasmid) is used as a negative control. All data were normalized to mode. **(B)** Table of representative binding patterns of the produced antibodies (also in 4, B and C) to different trypanosome cell lines. The (+) symbol indicates binding while the (-) non-binding. V and J segments as well as the CDR3 of both heavy and light are shown, along with the binding to the individual cell lines. The original isotypes and somatic hypermutations, if any, are also listed.

These results underscore the immunodominance of the relevant epitopes and the exquisite discrimination of the resulting antibodies. Specifically, they suggest that the immunodominant epitope that characterizes the VSG3_WT_ coat is limited to the immediate amino-acids surrounding the S317-*O*-glucose (including that sugar but excluding its companion on S319).

## Discussion

The antibody-mediated interactions between host and trypanosomes have been intensively studied over the years, but remain poorly understood. Investigating immune responses against the coat of the trypanosomes, together with the cognate VSG structures, will provide greater insights into the nature of such interactions. Here we show that mouse infection with trypanosomes of a specific VSG coat leads to robust plasma cell expansion, comprised of cells that are remarkably restricted in their antibody repertoire. This expansion is reminiscent of acute infection with Dengue in humans (Waickman et al., 2020). It is important to note however that repertoires elicited by Dengue are hypermutated and show a high degree of clonality, suggesting prior exposure. In contrast, repertoires elicited by VSGs are essentially unmutated (Fig. 4C and 6B). Furthermore, although each VSG gene appears to elicit a distinct V gene signature, the antibodies characterized by this signature are not clonal (i.e., they contain different D_H_ and J_H_ segments and consequently different CDR3 sequences). Though the existence of a robust serological response to trypanosome infection has been recognized for years (most recently by (Verdi and Zipkin et al., 2020), our work confirms that this response is polyclonal (Müller et al., 1996). Such polyclonal (and potentially autoreactive) antibodies have been documented in other infections as well, e.g., influenza (Lee et al., 2016, 2019; Woods et al., 2007), dengue (Correa et al., 2015; Waickman et al., 2020), hepatitis C (de Vita et al., 2008) and HIV (Moir and Fauci, 2009). Our work extends these findings by demonstrating that this polyclonal response is highly stereotyped between animals, is VSG-specific and is repertoire restricted.

The usage of distinct V gene signatures per VSG-elicited repertoire, which is highly reproducible between mice, implies the existence of common paratopes. In turn, this suggests that each VSG contains a small cohort of immunodominant epitopes to which the common paratopes might map. To date, co-crystal structures of VSGs and cognate antibodies are not available, and thus a definitive assignment of antibody-bound epitopes cannot be made. However, mutations in structurally-defined features of each VSG can lead to loss of antibody binding, suggesting that key immunodominant epitopes reside within a tightly circumscribed region of each VSG. For VSG2, such a feature is the calcium binding pocket, identified on the top lobe of the molecule: disruption of the pocket by point mutations in the calcium coordinating residues (DND) did not lead to structural alteration but did result in loss of recognition by VSG2_WT_ -elicited antisera and monoclonal antibodies. This implies that the subdomain of VSG2_WT_ that contains the DND motif forms an epitope that dominates the antibody response against this specific coat.

Similarly, for VSG3, an immunodominant epitope is defined by a small region surrounding the *O*-linked glucose on serine 317. Previous work had shown that this particular hexose anchors a chain of up to three additional hexoses to S317, leading Pinger et al (Pinger, Nešić and Ali, et al., 2018) to hypothesize that hexose heterogeneity contributed to repertoire diversification that would be detrimental to the immune response against the organism (Pinger, Nešić and Ali, et al., 2018). Indeed, we found that infections with VSG3_S317A_ coated trypanosomes elicited restricted antibody repertoires with a V_L_ gene signature dominated by V_L_ -gn33, but that this signature disappeared after infection with the wildtype VSG3 strain. This loss of signature correlates with increased virulence: infection with VSG3_S317A_ coated trypanosomes is naturally cleared while infection with the wildtype is lethal (Pinger et al., 2017). Perhaps equally interesting is the finding that a second hexose at a neighboring residue to S317 (S319) plays almost no role in defining the repertoire: mutating S319 to alanine resulted in a repertoire that matched the repertoire elicited by the wildtype VSG3 strain (Fig. 5C). Furthermore, mutating both serines resulted in a repertoire matching that elicited by the VSG3_S317A_ mutant (Fig. 5C). These findings illustrate the immunodominance of key epitopes on VSG3_WT_, which may be limited to the amino acids in the immediate vicinity of the *O*-glycan on S317, and suggest that the antibody repertoires elicited to specific VSGs not only respond to defined immunodominant features within each VSG but are highly discriminatory as they do so.

Overall, by examining two distinct VSGs and variants thereof, we demonstrate that the antibody response that mediates clearance to a given *T. brucei* coat can be hyperfocused to a restricted set of immunodominant epitopes that are surface exposed. Consequently, the immune response elicited by each *T. brucei* variant coat has the potential to be highly restricted and likely mediated by paratopes within the germline V regions of antibody genes. How *T. brucei* achieves this “immunofocusing” is likely related to the highly dense, repetitive and patterned nature of its surface coat. We hypothesize that this extreme focusing allows the parasite to use short-tract gene conversion or even point mutations (collectively referred to as mosaic formation (Hall et al., 2013; Mugnier et al., 2015) to expand its antigenicity well beyond what is encoded in its considerable genomic archive. By ensuring that cross reactivity between coat-defined epitopes will be exceedingly rare, even within similar variants, immunofocusing helps the parasite perpetually evade the immune response.

## Acknowledgments

We acknowledge synchrotron time at the Paul Scherrer Institut, Villigen, Switzerland (SLS, beamline X06DA, Vincent Olieric and colleagues), the Diamond Light Source (DLS, beamline i03, Neil Paterson and colleagues) and the Advanced Photon Source, Argonne National Laboratory (APS, beamline 24-ID-C, Jonathan P. Schuermann and colleagues). We thank Johan P. Zeelen for assistance with the synchrotron data acquisition and advice in structural determination. We thank Hedda Wardemann for assistance with B cell repertoire analyses and fruitful discussions especially during manuscript revision. We are grateful to Joey Verdi for beneficial comments and discussion during manuscript revision. Work in the F.N.P and C.E.S labs is funded by the Helmholtz Foundation. P.D.B.O and B.T.C are funded by National Institutes of Health P41 GM109824 and P41 GM103314 to B.T.C. R.B and T.R. are supported by the German Research Foundation to T. R. (Ru 747/1-1) within in the framework of FOR 2509.

## Author contributions

Conceptualization, Resources and Funding acquisition: F.N.P., C.E.S., Investigation: A.G., F.A.B., G.T., E.P.V., M.L., C.E.S., Mass Spectrometry (MS) analysis: P.D.B.O., B.T.C., R.B., T.R., ITC analysis: K.P., Formal Analysis: A.G., F.A.B., C.E.S., P.D.B.O.(MS), B.T.C.(MS), K.P.(ITC), R.B.(MS), T.R.(MS), Visualization: A.G., F.A.B., C.E.S., P.D.B.O.(MS), B.T.C.(MS), K.P.(ITC), R.B.(MS), T.R.(MS) Writing – Initial manuscript: F.N.P., C.E.S., Writing – Revised manuscript: F.N.P., C.E.S., A.G., F.A.B., G.T.

## Lead Contact

Further information and requests for resources and reagents should be directed to and will be fulfilled by the lead contact, Nina Papavasiliou.

## Data availability

Structures that support the findings of this study have been deposited in RCSB Protein Data Bank with IDs: 7P56 (VSG2_WT_), 7P57 (VSG2_AAA_), 7P59 (VSG3_WT_), 7P5A (VSG3_S317A_), 7P5B (VSG3_S319A_), 7P5D (VSG3_SSAA_); Mass spectrometry data are available via ProteomeXchange with identifier PXD027384; Full repertoire data are available from the corresponding authors upon reasonable request. All other data is available in the main text or the supplementary data.

## Star Methods

### Trypanosome cell lines and mouse strains

All trypanosome cell lines used in this study were bloodstream-form trypanosomes derived from the Lister-427 “2T1” cell line (Alsford et al., 2005) and were cultivated *in vitro* in HMI-9 medium (Hirumi and Hirumi, 1989) (formulated as described by PAN Biotech without FBS, l-cysteine or β-mercaptoethanol), supplemented with 10% fetal calf serum (Gibco), l-cysteine and β-mercaptoethanol. Cells were cultured at 37 °C and 5% CO2. The VSG3_WT_ and VSG3_S317A_ cell lines were described before (Pinger, Nešić and Ali, et al., 2018). Isogenic VSG3_S319A_ and VSG3_SSAA_ mutant clones were derived from transfection of 2T1 (VSG2 expressing) cells with the pAG plasmids described below, and then initially selected based on loss of VSG2 expression and gain of VSG3 expression (by FACS, see below). Derivation of isogenic VSG2_WT_ or VSG2_AAA_ mutant trypanosomes was achieved through transfection of naturally occurring switchers of 2T1 (expressing VSG3, or VSG9) with the relevant pFAB plasmids (see below), and then initially selected based on gain of VSG2 expression.

These strains were used to infect female, wild-type C57BL/6J mice, aged 6–8 weeks at experiment start (Janvier). Mice were kept in IVC cages under SPF conditions in the animal facility at the German Cancer Research Center (DKFZ, Heidelberg). All animal experiments were performed in accordance with institutional and governmental regulations, and were approved by the Regierungspräsidium (Karlsruhe, Germany).

### Engineering plasmids used to generate isogenic *T. brucei* clones expressing VSG2 or VSG3 and mutants thereof

Plasmids used to generate VSG3_WT_ and VSG3_S317A_ -expressing cells were described in (Pinger, Nešić and Ali, et al., 2018). Modifications of these plasmids using site-directed mutagenesis (Q5, New England Biolabs) were used to generate VSG3_S319A_ (pAG1) and the VSG3_SSAA_ -expressing cells (pAG2). Plasmids were first linearized by EcoRV (New England Biolabs), and then transfected into VSG2-expressing cells (2T1).

Plasmids used to generate VSG2 and VSG2_AAA_ isogenic strains (pFAB1 and pFAB4 respectively) were created using vector pSY371D-CTR-BSD (Pinger et al., 2017) as a backbone (also modified to contain the hygromycin-resistance gene (HYG) instead of blasticidin). pFAB1 and pFAB4 were linearized with BglII (New England Biolabs) prior to transfection.

### *T. brucei* transfections, clone selection and verification

All transfections were performed using 10ug of each plasmid mixed with 100ul of 2.5-3x10^7^ cells in homemade Tb-BSF buffer (90mM Na_2_HPO_4_, pH 7.3, 5mM KCl, 0.15mM CaCl_2_, 50mM HEPES, pH7.3), using the AMAXA nucleofector (Lonza) program X-001, as previously described (Burkard et al., 2007). After 6h, blasticidin at a concentration of 100ug/ml (VSG3_S319A_ and VSG3_SSAA_ cell lines) or hygromycin at 25ug/ml (VSG2_WT_ and VSG2_AAA_ cell lines) were added and single-cell clones were obtained by serial dilutions in 24-well plates and harvested after 5 days. The VSG3_S319A_ and VSG3_SSAA_ clones were screened by FACS for VSG3 expression and VSG2 loss of expression and VSG2_WT_ and VSG2_AAA_ clones for VSG2 expression using monoclonal antibodies against these VSGs (Pinger et al., 2017). Positive clones were sequenced by isolating RNA using the RNeasy Mini Kit (Qiagen), followed by DNAse treatment with the TURBO DNA-free kit (Invitrogen) and cDNA synthesis with ProtoScript II First Strand cDNA Synthesis (New England Biolabs). The sequences were then amplified, using Phusion High-Fidelity DNA Polymerase (New England Biolabs), a forward primer binding to the spliced leader sequence and a reverse binding to the VSG 3′untranslated region. The final products were purified by gel extraction from a 1% gel with the NucleoSpin Gel and PCR clean-up kit (Macherey-Nagel) and sent for Sanger sequencing.

### Spleen and antisera collection

6-8-week-old C57BL/6J wild-type mice were injected intraperitoneal (i.p.) with 1 × 10^3^ parasites in HMI-9. Mice that cleared VSG3_S317A_ infections naturally, were injected i.p. with 100 parasites in HMI-9. Mice treated with diminazene underwent an injection (i.p.) of 250 ng diminazene aceturate (Abcam)/mouse 4 days after infection, and this procedure was repeated after 24 h. On day 8 post parasite injection, mice were euthanized with CO2. Mice that naturally cleared the first peak of parasitemia were carefully monitored three times per day between days 5-7 post infection and euthanized with CO2 upon clearance of parasitemia.

After mice were euthanized, blood was collected via cardiac puncture and serum was separated from whole blood using Microtainer SST serum collection tubes (BD 365968). Spleen was also collected and single-cell suspensions were prepared by standard procedures.

### Flow Cytometry and Single Cell Sorting

Splenocytes from spleens of Trypanosome-infected mice were thawed, washed in RPMI media (Gibco) at room temperature, centrifuged at 2000 rpm for 5 min and resuspended in 100ul 2% FBS/PBS. Cells were stained with rat anti-mouse CD19-BV421 (1:100, Biolegend), rat anti-mouse CD138-BV510 (1:300, Biolegend), rat anti-mouse IgG1-BV650 (1:100, Biolegend) and goat anti-mouse IgM-Biotin (1:400, Jackson Laboratories) for 45 min on ice in the dark. Biotin was detected using Streptavidin-BV785 (1:400, Biolegend) and 7-Aminoactinomycin D (7AAD) (1:200, Invitrogen) was used as a dead cell marker in all samples. The cells were analyzed on a LSRFortessa instrument (BD Bioscience), single-cell sorted into 384-well plates (black frame, 4titude) using either an Aria I or Aria Fusion II cell sorter (BD Bioscience) and analyzed using FlowJo software (v10). For single-cell sorting, the plasma cell population was defined as 7AA^-^CD19^lo^CD138^+^ and was checked for IgM and IgG1 surface expression, without including these markers in the gating of the sort population. The exact isotype of each plasma cell was determined later by sequence analysis.

### Antibody repertoires, cloning and production of mouse antibodies

Ig heavy (IgH), kappa (Igκ) and lambda (Igλ) genes from single plasma cells were amplified as described before (Tiller et al., 2008). Briefly, cDNA of each cell was produced using random hexameric primers and then the IgH, Igκ and Igλ gene transcripts were amplified by a subsequent semi-nested PCR approach using V segment and constant region-specific primers (Tiller et al., 2008). The PCR products were sent for Sanger sequencing and analyzed with the IgBlast online tool (NCBI, (Ye et al., 2013)) to generate the antibody repertoires. Matching heavy and light pairs originating from the same cell were picked, amplified using V and J gene-specific primers that include restriction sites for cloning and cloned into appropriate human expressing vectors (Tiller et al., 2008). After they were successfully cloned and their sequence was validated, the corresponding heavy and light chain plasmids were co-transfected into adherent HEK293T cells (ATCC) using Lipofectamine 2000 (Invitrogen) reagent in 1x OptiMEM (Gibco). After 6h the media was removed and cells were supplemented with 1x Nutridoma-SP (Roche). Supernatants containing the produced antibodies (as IgG1) were collected after 48h and 96h and tested for their ability to bind to live trypanosomes by FACS.

### VSG purification

VSGs were purified as described in (Cross, 1984) with minor changes. Briefly, cells grown to a density of 2.5-4 × 10^6^ were centrifuged and lysed in 0.2mM ZnCl_2_. Following cell lysis, the mixture was centrifuged and the pellet containing the VSG protein was resuspended in prewarmed (40°C) 20mM HEPES buffer, pH 8.0, supplemented with 150mM sodium chloride (VSG2_WT_ and VSG2_AAA_) or 20mM HEPES buffer, pH 7.5, supplemented with 150mM sodium chloride (VSG3_WT_ and the three sugar-mutants). After a second centrifugation, the supernatants containing the VSG protein were loaded onto an anion-exchange column (Q Sepharose Fast Flow, GE Healthcare) previously equilibrated with the respective HEPES buffer as above. The flow-through and two washes containing the VSG of interest were collected and concentrated using an Amicon Stirred Cell (Merck Millipore) and the sample was then run over a gel filtration column (Superdex 200, GE Healthcare) after equilibration with the respective HEPES buffer as above. Aliquots of both the different purification steps and the gel filtration runs were subjected to SDS-PAGE analysis for visual inspection (Figure S3, C-H). From the gel filtration step onwards, all VSG3 constructs were gradually carboxy-terminal (CTD) truncated, likely due to cleavage by endogenous proteases, resulting in the crystallization of only the N-terminal domain. For crystallization of VSG2_AAA_, VSG purified from gel filtration was concentrated to 2.2mg/ml and left for 1 week at 4°C where the CTD was also truncated (again, likely due to endogenous proteases). VSG2_AAA_ underwent a second gel filtration (Superdex 200, GE Healthcare) and was concentrated to 4.8mg/ml. For crystallization of VSG2_WT_, VSG was first purified with the same protocol but using 20mM Tris, pH 8.0. After passage over an anion-exchange column, VSG was concentrated to 10mg/ml and passed over a gel filtration column (Superdex 200, GE Healthcare) equilibrated in 20 mM Tris pH 8.0. Aliquots from the gel filtration runs were subjected to SDS–PAGE analysis for visual inspection, and concentrated to 2.5mg/ml.

### Crystallization and structural determination

Native crystals of VSG2_WT_ were grown by vapor diffusion using hanging drops formed from mixing a 1:1 volume ratio of the protein with an equilibration buffer consisting of 0.1M K-acetate, 22% PEG8000. For cryoprotection, crystals were transferred directly into a buffer with a 25% PEG 8000, 0.1M K-acetate, 20% glycerol and flash-cooled immediately afterward to 100 K (−173.15 °C). Data were collected at Advanced Photon Source (APS) at Argonne National Laboratory at beamline 24-ID-C and processed onsite through the RAPD software pipeline. The structure was solved by molecular replacement using the model of PDB entry 1VSG with the PHENIX software suite (Adams et al., 2010). The model was improved and finalized through several cycles of auto-building (PHENIX), manual adjustment, and refinement (PHENIX) (Table S1).

Native crystals of VSG2_AAA_ were grown by vapor diffusion using hanging drops formed from mixing a 1:1 volume ratio of the protein with an equilibration buffer consisting of 0.1M Tris pH 8.0, 39% PEG400. This equilibration buffer is cryoprotective and crystals were flash-cooled afterward to 100 K (−173.15 °C). Data were collected at the Swiss Light Source (SLS) at a wavelength of 1.0Å on beamline X06DA (PXIII) The structure was solved by molecular replacement using the model of PDB entry 1VSG with the PHENIX suite (Adams et al., 2010). The model was improved and finalized through several cycles of auto-building (PHENIX), manual adjustment, and refinement (PHENIX) (Table S1).

Purified VSG3_WT_ and the three sugar-mutants were concentrated to 2mg/ml in 20mM HEPES buffer, pH 7.5, supplemented with 150mM NaCl. Crystals were grown at 22°C by vapor diffusion using hanging drops with a 1:1 volume ratio of protein to equilibration buffer consisting of 21% PEG 3350, 250mM NaCl and 100mM Tris, pH 8.2 for VSG3_WT_ and VSG3_S317A_ and 25% PEG 3350, 300mM NaCl and 100mM HEPES, pH 7.5 for VSG3_S319A_ and VSG3_SSAA_. For cryoprotection the crystals were transferred to the same buffer as that used for equilibration but supplemented with 25% v/v glycerol and were flash-frozen in liquid nitrogen. Data for VSG3_WT_, VSG3_S317A_ and VSG3_SSAA_ were collected at the Swiss Light Source (SLS) at a wavelength of 1.0Å on beamline X06DA (PXIII) and for VSG3_S319A_ at the Diamond Light Source at a wavelength of 0.9763Å on beamline i03. The VSG3_WT_ and sugar-mutant structures were obtained using the previously solved VSG3_WT_ structure (PDB ID: 6ELC) (Pinger, Nešić and Ali, et al., 2018) as a model to perform Molecular Replacement in the PHENIX suite (Adams et al., 2010). The models were improved and finalized through several cycles of auto-building (PHENIX), manual adjustment, and refinement (PHENIX) (Table S2).

### Mass Spectrometry

Purified VSG2_WT_ and VSG2_AAA_ samples were buffer-exchanged into native Mass Spectrometry (nMS) solution (150mM ammonium acetate, 0.01% Tween-20, pH7.5) using Zeba desalting microspin columns with a 40-kDa molecular weight cut-off (Thermo Scientific). An aliquot (2–3µL) of the buffer-exchanged sample was loaded into a gold-coated quartz capillary tip that was prepared in-house. The sample was then electrosprayed into an Exactive Plus EMR instrument (Thermo Fisher Scientific) using a modified static nanospray source (Olinares and Chait, 2020). The MS parameters used included: spray voltage, 1.1 – 1.3kV; capillary temperature, 200°C; S-lens RF level, 200; resolving power, 35,000 at *m/z* of 200; AGC target, 1 – 3 x 10^6^; number of microscans, 5; maximum injection time, 200ms; in-source dissociation (ISD), 200V; injection flatapole, 8V; interflatapole, 7V; bent flatapole, 6V; high energy collision dissociation (HCD), 10V; ultrahigh vacuum pressure, 5 × 10^−10^ mbar; total number of scans, 100. Mass calibration in positive mode was performed using cesium iodide. Raw nMS spectra were visualized using Thermo Xcalibur Qual Browser (version 4.2.47). Deconvolution was performed using UniDec version 4.2 (Marty et al., 2015; Reid et al., 2019). The UniDec parameters used were m/z range: 3,500 – 5,500; mass range: 90,000 – 105,000Da; sample mass every 0.2Da; smooth charge state distribution, on; peak shape function, Gaussian; and Beta softmax function setting, 20.

Native MS analyses of the purified VSG2 samples revealed massive heterogeneity due to extensive glycosylation consistent with previous studies that determined N-glycosylation at two Asn sites and variable galactosylation at the C-terminal GPI anchor per VSG2 monomer (Manthri et al., 2008; Mehlert et al., 1998; Zamze et al., 1991). To reduce sample complexity and enable unambiguous identification of bound metal ions, the VSG samples were N-deglycosylated using PNGaseF (NEB) with and without 10mM EDTA at 37°C for 3 - 4h. The deglycosylated samples were then buffer exchanged and analyzed by nMS as described above.

The expected masses for the VSG2 glycoforms were determined as follows. The signal peptide at the N-terminus (residues 1-27) and the GPI attachment signal peptide at the C-terminus (residues 460-476) were removed from the precursor VSG2_WT_ sequence yielding a monomer mass 46,291.8Da for the processed protein. Common to all the monomers is the presence of four disulfide bridges (-8.06 Da) as well as attachment of glucosamine-α1-6-myo-inositol-1,2-cyclic phosphate (+385.26 Da), ethanolamine phosphate (+123.05 Da) and five hexoses (+810.70 Da) at the C-terminal GPI anchor. Moreover, PNGaseF cleavage of N-linked glycans at two Asn residues results in deamidation of Asn to Asp (+1.97 Da). Overall, the resulting monomer and dimer masses are 47,604.7 Da and 95,209.4 Da, respectively. Each additional galactose attachment from variable galactosylation at the GPI anchor adds 162.14 Da. Each Ca^2+^ ion bound adds 38.06 Da (average mass of Ca minus the mass of 2H^+^ from deprotonation of two aspartic acids that coordinate with the cation). The DND-to-AAA mutation results in a decrease of VSG2_WT_ dimer mass by 262.10 Da (Table S3).

Purified VSG3_WT_ was either concentrated to 3mg/mL in 20mM HEPES, pH 8.0, with 150mM NaCl and sent for Electron transfer dissociation (ETD) analysis or 50ug of protein in the same buffer were treated as described in (Pinger, Nešić and Ali, et al., 2018) and then separated by SDS PAGE. The 17 kDa fragment was excised and processed as described (Fecher-Trost et al., 2013). In brief, trypsin digestion was done overnight at 37°C. The reaction was quenched by addition of 20µL of 0.1% trifluoroacetic acid (TFA; Biosolve, Valkenswaard, The Netherlands) and the supernatant was dried in a vacuum concentrator before LC-MS analysis. Nanoflow LC-MS^2^ analysis was performed with an Ultimate 3000 liquid chromatography system coupled to an Orbitrap Elite mass spectrometer equipped with ETD (Thermo-Fischer, Bremen, Germany). Samples were dissolved in 0.1% TFA, injected to a self-packed analytical column (75um x 200mm; ReproSil Pur 120 C18-AQ; Dr Maisch GmbH) and eluted with a flow rate of 300 nl / min in an acetonitrile-gradient (3% - 40%). The mass spectrometer was operated in data-dependent acquisition mode, automatically switching between MS and MS^2^. Collision induced dissociation MS^2^ spectra were generated for up to 10 precursors with normalized collision energy of 29%. Electron transfer dissociation (ETD) MS^2^ spectra were generated for up to 5 precursors using the default settings of the instrument. Each analysis was done in triplicate.

Raw files were processed using Proteome discoverer 2.2 (Thermo Scientific) for peptide identification and quantification. MS^2^ spectra were searched against the Uniprot Trypanosoma database (UniprotKB), a custom database entry with the VSG3 sequence and the contaminants database (MaxQuant database; MPI Martinsried) with the following parameters: Acetyl (Protein N-term), Oxidation (M) and Hex (S,T) as variable modifications and carbamidomethyl (C) as static modification. Trypsin/P was set as the proteolytic enzyme with up to 2 missed cleavages allowed. The maximum false discovery rate for proteins and peptides was 0.01 and a minimum peptide length of 7 amino acids was required.

### Isothermal Titration Calorimetry

To evaluate binding of ions to VSG2_WT_ or VSG2_AAA_ and remove contaminating calcium, all glassware was first treated with phosphoric acid and water to prepare all buffers and solutions for treatment with Chelex 100 Resin 200-400 mesh (BioRad). Upon purification of VSG2, the CTD was removed by digestion with Endoproteinase LysC (New England Biolabs) at a 1:500 LysC/substrate ratio by mass together with 10mM EDTA for 5h at 37°C. The samples were then run over a gel filtration column (Superdex 200, GE Healthcare) after equilibration with the respective 20mM Tris pH 8.0 buffer. Afterwards, proteins were concentrated to 30mM by concentration in 10-kDa disposable ultrafiltration centrifugal devices.

ITC experiments were performed using a PEAQ ITC (Malvern) at 20°C with a stirring rate of 750 rpm. Titration buffers contained 20mM Tris pH 8. In each experiment, the protein concentration in the cell was 30µM. CaCl_2_, KCl and MgCl_2_ were injected at 600µM with injection sizes of 12×3µl. The data were baseline-corrected, integrated and analysed with the PEAQ ITC Analysis software (Malvern), fitted using a single-site binding model.

### Fluorescence-activated cell sorting (FACS)

To assess binding of antisera to live trypanosomes, 1 x 10^6^ parasites were harvested and incubated with VSG2_WT_ anti-sera (1:4000) or VSG2_AAA_ (1:2000) together with Fc block (1:200, BD Pharmingen) in cold Trypanosome Dilution Buffer (TDB) (5mM KCl, 80mM NaCl, 1mM MgSO_4_, 20mM Na_2_HPO_4_, 2mM NaH_2_PO_4_, 20mM glucose, pH 7.4) for 10 min at 4°C. Cells were washed once with cold TDB and resuspended in 200µl cold TDB with rat anti-mouse IgM-FITC (1:500, Biolegend). Upon one wash with cold TDB, cells were resuspended in 250µl TDB and immediately analyzed with FACSCalibur (BD Bioscences) and FlowJo software (v10).

To verify whether the repertoire antibodies that were produced in HEK cells were able to bind to live trypanosomes, 0.5 x 10^6^ parasites were harvested, washed once with cold HMI-9 without FBS and stained in 200ul of each antibody supernatant for 10 min at 4°C. Cells were pelleted, resuspended in 100ul cold HMI-9 without FBS with mouse anti-human IgG1-AlexFluor488 (1:500, Invitrogen) for 10 min at 4°C in the dark. Cells were washed once with cold HMI-9 without FBS, resuspended in 100ul of the same buffer and immediately analyzed with FACSCalibur (BD Bioscences) and FlowJo software (v10).

## Supplementary Information

Figures S1-S7 Tables S1-S3

