## Supplementary figures, methods, and tables for "Immunodominant surface epitopes power immune evasion in the African trypanosome"

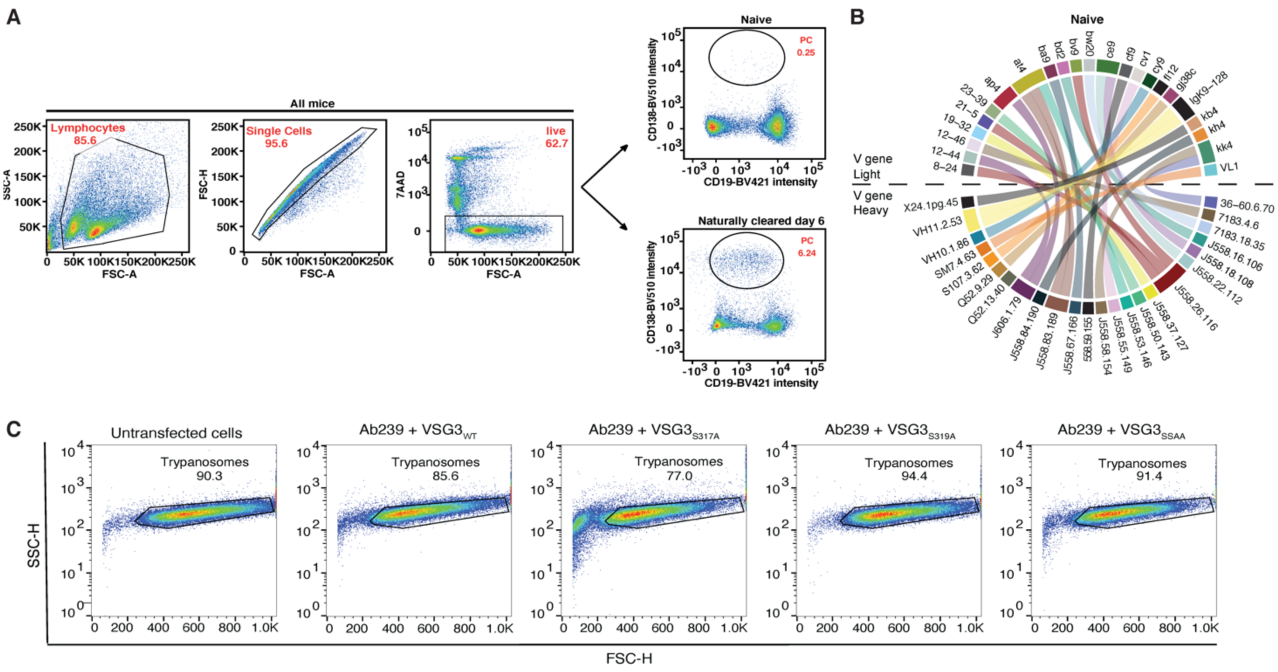

**Figure S1: Overall gating strategies for naïve and infected mice, together with the plasma cell antibody repertoire of a naïve mouse. (A)** Gating strategy for the sorting of plasma cells from naïve or infected mice. The plots shown are for infections with VSG2 coated parasites but identical plots were collected for infections with VSG3 coated trypanosomes and their variants **(B)** Circos plot generated for the antibody repertoire of one naive mouse (n=29 pairs, V gene signatures shown only). Different colors depict each heavy chain variable gene (bottom half of the plot), and each light chain variable gene (top half of the plot).  The heavy and light chain variable gene pairings that form the antibodies are illustrated as connector lines. Shannon entropy shows clonal diversity on a sequence level, with a value of 1.0 representing 100% clonal diversity (no clones), while a value of 0.0 corresponding to 0% clonal diversity (only clones). **(C)** Gating strategy for each VSG3 cell line stained with recombinant Ab239. The "Trypanosomes" gate was used for data acquisition and analysis, for which 25.000 events were recorded. The same methodology and gating were applied to the other FACS experiments with monoclonal antibodies and polyclonal antisera (e.g. Figures 2E, 2F, 3C and 4D and Supplementary Information Figures S2, S7 and S8).

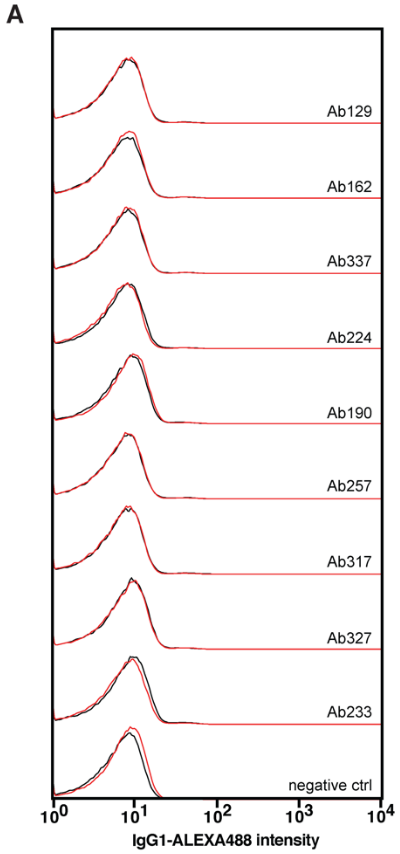

B

| Ab | IgHV | CDR3-H | IgLV | CDR3-L | Binding | Isotype | Mutations |
|---|---|---|---|---|---|---|---|
| 129 | J558.49.141 | ARGHYDYDGGAMDY | VL1 | ALWYSNHLV | - | IgG2b | 0 |
| 162 | J558.49.141 | ARNYGNYGYYAMDY | VL1 | ALWYSNHWV | - | IgM | 0 |
| 337 | J558.49.141 | ARAYDYDGDYYAMDY | VL1 | ALWYSNHWV | - | IgM | 0 |
| 224 | J558.49.141 | ARSGNYYGSSRDYYAMDY | VL1 | ALWYSNHWV | - | IgM | 0 |
| 190 | J558.49.141 | ARAYDYDGDYYAMDY | VL1 | ALWYSNHWV | - | IgM | 0 |
| 257 | J558.49.141 | ARGAYYSNLYAMDY | VL1 | ALWYSNHFI | - | IgM | 0 |
| 317 | J558.49.141 | ARALYGSSSFDY | VL1 | ALWYSNHWV | - | IgM | 0 |
| 327 | J558.49.141 | ARDYYSNLYAMDY | VL1 | ALWYSNHFI | - | IgM | 0 |
| 233 | J558.49.141 | ARPSSYAAEDYYAMDY | VL1 | ALWYSNHWV | - | IgM | 0 |

**Figure S2: VSG2$_{WT}$ elicited antibodies that lack the dominant heavy chain V gene signature (V*H*10) do not bind to live trypanosomes. (A)** FACS histograms showing binding intensities of recombinant antibodies with wild type parasites (black) and AAA-mutant parasites (red). These antibodies represent non-dominant V gene signatures (vs Figure 1C-D). Staining with supernatants from untransfected cells (no plasmid) is used as a negative control. All data are normalized to mode. **(B)** Table with the antibodies presented in panel a, showing the heavy and light chain V genes. The (-) symbol indicates no binding. The CDR3s from each antibody is shown along with the original antibody isotype and somatic hypermutations.

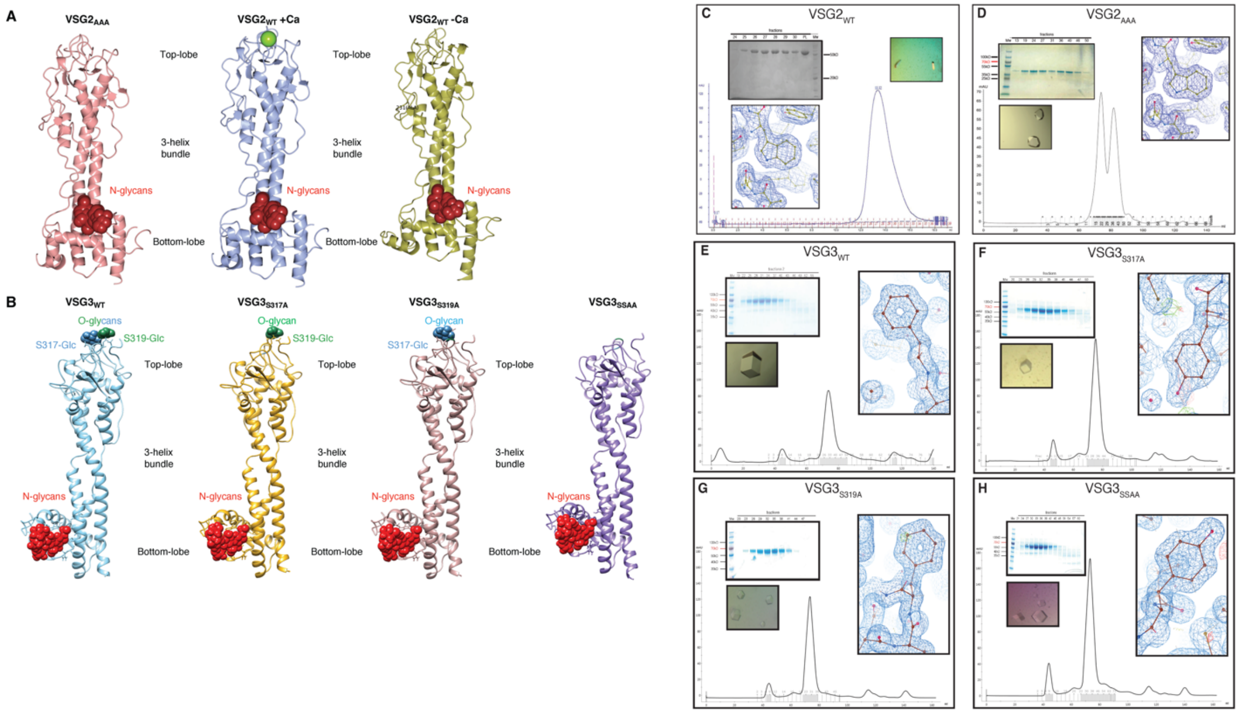

**Figure S3: Crystal structures of VSG2 and VSG3 and crystal information. (A)** The crystal structures of VSG2$_{AAA}$ (pink), VSG2$_{WT}$ with calcium (cyan) and VSG2$_{WT}$ without calcium (gold) shown side by side as ribbon diagrams (only a single monomer of VSG2 is shown). The N-glycans are represented as red spheres on the bottom lobe and the calcium ion as a green sphere. **(B)** The crystal of VSG3$_{WT}$ shown here as a ribbon diagram colored in light blue. The N-glycans are represented as red spheres on the bottom lobe, while the *O*-linked sugars as blue (S317-Glc) and green (S319-Glc) spheres on the top lobe. The crystals of VSG3$_{S317A}$, VSG3$_{S319A}$ and VSG3$_{SSAA}$ are also shown here in gold, soft brown and purple respectively. The same parameters as for VSG3$_{WT}$ apply for these three as well. **(C)** The panels show a gel filtration chromatogram (bottom) of purified VSG2$_{WT}$, a representative coomassie stained SDS-PAGE gel of the fractions after gel filtration (top left), a VSG2$_{WT}$ crystal before collection and finally, a 2Fo-Fc electron density contoured at 1σ, after final refinement. Similar panels can be seen for the VSG2$_{AAA}$ protein in **(D)**. **(E)** The panels show a gel filtration chromatogram (bottom) of purified VSG3$_{WT}$, a representative coomassie stained SDS-PAGE gel of the fractions after gel filtration (top left), a VSG3$_{WT}$ crystal before collection (bottom left) and finally, a 2Fo-Fc electron density contoured at 1σ, after final refinement. Similar panels can be seen for the other proteins: **(F)** VSG3$_{S317A}$, **(G)** VSG3$_{S319A}$ and **(H)** VSG3$_{SSAA}$.

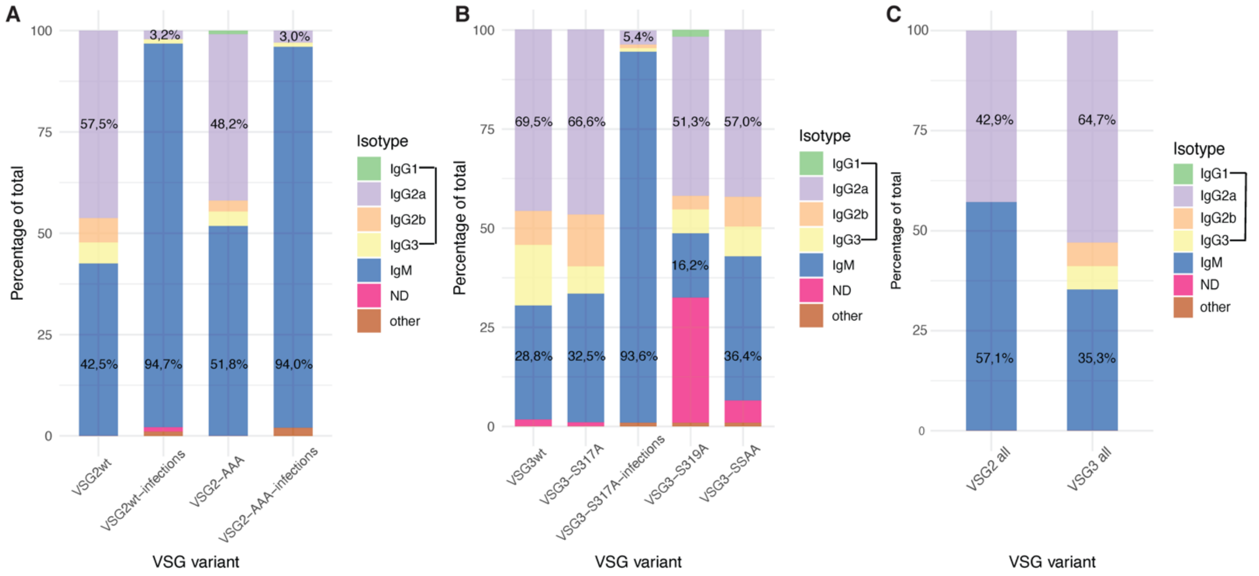

**Figure S4: Isotype distribution for antibodies elicited by VSG2 and VSG3.** Original isotypes from the VSG2 and mutant **(A)** and VSG3 and mutant(s) **(B)** repertoires, from both diminazene-treated and naturally-cleared infection experiments. The percentages of IgM (blue) and IgGs (IgG1-green, IgG2a-purple, IgG2b – orange and IgG3– yellow) can be seen on the plot. ND stands for „Non-Determined“ (in fuchsia). The x-axis represents the different variants, while the y-axis represents the relevant percentages (up to 100%). Within each bar, the exact percentages for IgGs and IgMs are shown. The IgG percentage includes all sub-classes (IgG1, IgG2a, IgG2b and IgG3). **(C)** Isotypes of the antibodies that, as recombinants, successfully bound to the cognate cell lines, raised against all VSG2 variants (left bar) or VSG3 variants (right bar).

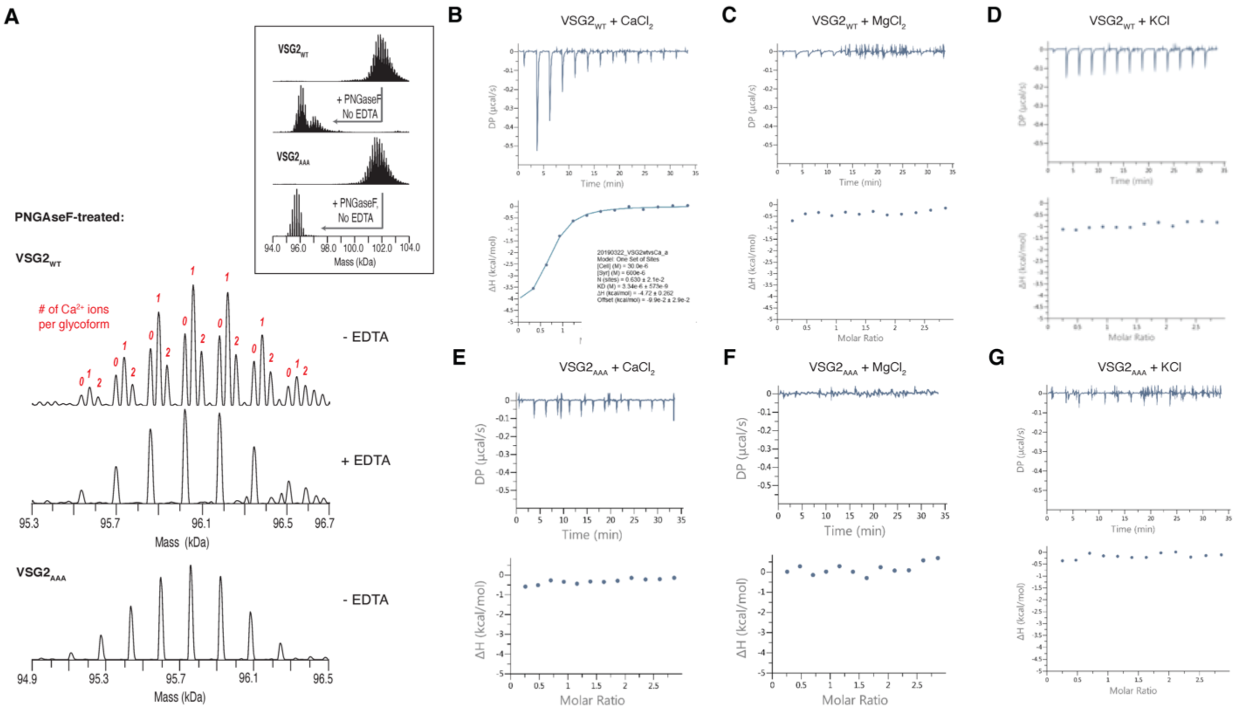

**Figure S5: VSG2$_{WT}$ binds calcium. (A)** Deconvolved spectra from high-resolution native MS analyses of VSG2 samples. PNGaseF treatment removed the N-linked glycans reducing the heterogeneity and mass of VSG2 dimers by at least 5 kDa (inset). The remaining glycoform microheterogeneity comes from variable galactosylation at the C-terminal GPI anchor (up to 8 Gal molecules per VSG2 dimer, see Extended Data Table 3). N-deglycosylation of VSG2$_{WT}$ revealed triplet peaks spaced 38 – 40 Da apart corresponding to the coordination of zero, one or two $Ca^{2+}$ ions for each glycoform (Table S3). Addition of EDTA, a chelating agent during PNGaseF incubation of VSG2$_{WT}$ completely removed the coordinated $Ca^{2+}$ ions. In contrast, no $Ca^{2+}$-bound glycoforms were observed with PNGaseF treatment of VSG2$_{AAA}$ under EDTA-free conditions. **(B-G)** ITC data for $CaCl_2$, $MgCl_2$ and KCl binding to each VSG2 protein. The upper panels contain the baseline corrected raw data, and the lower panel contains the peak-integrated, concentration normalized data for the heat of reaction vs. molar ratio of each individual metal per VSG2 protein. **(B)** VSG2$_{WT}$ was measured with two independent replicates: 300µM $CaCl_2$ was titrated into 30µM VSG2$_{WT}$, the curve fitted with a single binding site model to calculate a Kd of 3.92 +/- 0.58 nM and N of 0.645 +/- 0.015. **(C)** VSG2$_{WT}$ was measured once using the same conditions as in (B) but with $MgCl_2$. No binding was detected. **(D)** VSG2$_{WT}$ was measured once using the same conditions as in (B) but with KCl. No binding was detected **(E)** VSG2$_{AAA}$ was measured with two independent replicates: 300μM $CaCl_2$ was titrated into 30μM $VSG2_{AAA}$. No binding was detected. **(F,G)** similar measurements with $MgCl_2$ or KCl show no detectable binding.

A

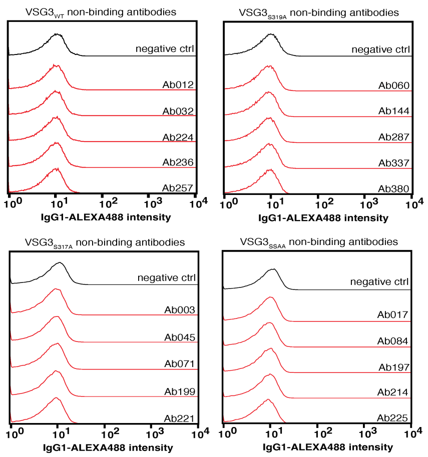

B

| Ab | IgkV | IgkJ | CDR3-k | IgHV | CDR3-H | Binding | Isotype | Mutations |
|---|---|---|---|---|---|---|---|---|
| 012 | 21-5 | JK4 | QQSNEDPFT | J558.67.166 | ARDYYGTSFAY | - | IgG2a | 0 |
| 032 | 21-4 | JK1 | QQSNEDPPT | 36-60.8.74 | ARAASYYYGSSYWYFDV | - | IgG2b | 0 |
| 224 | kk4 | JK5 | QQWSSNPPT | 3609.7.153 | ARIVWDGYYVTY | - | IgG3 | 0 |
| 236 | 21-10 | JK5 | QQNNEDPLT | 36-60.8.74 | ARNSPYYYGSSRVFDV | - | IgG2a | 0 |
| 257 | bt20 | JK4 | LQSDNLPLT | J558.75.177 | ARRGLTGPFDY | - | IgG2a | 0 |
| 060 | gn33 | JK5 | QQYWSTPLT | 3609N.2.77 | SIDYDGLGYFDV | - | IgM | 0 |
| 144 | gr32 | JK1 | QQGQSYPLT | J558.18.108 | ARGYYGSTYYYAMDY | - | IgG2a | 0 |
| 287 | ce9 | JK4 | QQGNTLFT | J558.72.173 | ARFYYGSSYFDY | - | IgG2a | 0 |
| 337 | ce9 | JK1 | QQGNTLPWT | 3609.7.153 | ARIPIYDGLYYAMDY | - | IgG3 | 0 |
| 380 | ce9 | JK2 | QQGNTLYT | 3609.7.153 | ARLLLNGYYAMDY | - | IgM | 0 |
| 003 | bt20 | JK2 | LQSDNLPYT | J558.85.191 | ASRYGSSPDFAY | - | IgG2a | 0 |
| 045 | 21-5 | JK5 | QQSNEDPLT | J558.6.96 | ARKGDYDWYFDV | - | IgM | 0 |
| 071 | 21-2 | JK2 | QQSKEVPYT | J558.55.149 | ARSHSSGYVGAMDY | - | IgM | 0 |
| 199 | bd2 | JK1 | WQGTHFPWT | J558.84.190 | AREGDYYAMDY | - | IgG2a | 0 |
| 221 | gn33 | JK2 | QQYWSTPYT | 36-60.6.70 | ARGGLGSY | - | IgG2a | 0 |
| 017 | ci12 | JK4 | QQLYSTPLT | J558.6.96 | ARGSSGYDQYYFDY | - | IgM | 0 |
| 084 | gn33 | JK2 | QQYWSTPYT | 36-60.6.70 | ARDGGLP | - | IgM | 0 |
| 197 | ci12 | JK1 | QQLYSTPLT | J558.6.96 | ARKGVYYDYDGYFDY | - | IgM | 0 |
| 214 | ci12 | JK5 | QQLYSTPLT | J558.26.116 | ARRGYYDYDVGYY | - | IgM | 0 |
| 225 | bt20 | JK4 | LQSDNLPLT | 36-60.4.66 | ARGSSGYAMDY | - | IgM | 0 |

C

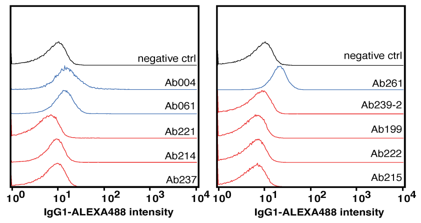

D

| Ab | IgLV | IgLJ | CDR3-L | IgHV | CDR3-H | Binding S317A | Isotype | Mutations |
|---|---|---|---|---|---|---|---|---|
| 004 | gn33 | JK2 | QQYWSTPYT | J558.26.116 | ARKGLHYWYFDV | + | IgM | 0 |
| 061 | gn33 | JK2 | QQYWSTPYT | 36-60.6.70 | ASYGYDVGWFAY | + | IgG2a | 0 |
| 221 | gn33 | JK2 | QQYWSTPYT | 36-60.6.70 | ARGGLGSY | - | IgG2a | 0 |
| 214 | gn33 | JK2 | QQYWSTPYT | 36-60.6.70 | ARGGSSGYDY | - | IgG2a | 0 |
| 237 | gn33 | JK2 | QQYWSTPYT | 7183.20.37 | ARGGYEFLYYFDY | - | IgM | 1 |
| 261 | gn33 | JK4 | QQYWSTPFT | J558.67.166 | ARRGVVDYFDY | + | IgM | 0 |
| 239-2 | gn33 | JK4 | QQYWSTPFT | J558.55.149 | ASTLYGNYEGRFAY | - | IgM | 0 |
| 199 | gn33 | JK4 | QQYWSTPFT | 36-60.6.70 | ARDSSGYGGAY | - | IgG2a | 1 |
| 222 | gn33 | JK4 | QQYWSTPFT | J558.53.146 | ATYGSSYVGYFDV | - | IgM | 0 |
| 215 | gn33 | JK4 | QQYWSTPFT | VH7183.5b | ARPLIYDGLFAY | - | IgG2b | 0 |

**Figure S6: Representative VSG3$_{WT}$ – and VSG3-sugar-mutant elicited antibodies and VSG3$_{S317A}$– elicited gn33 antibodies that pair with different heavy chains, that are not able to bind to cognate trypanosomes. (A)** Histograms showing lack of binding of a few selected antibodies to VSG3$_{WT}$ and mutant cell lines as indicated by the labeling. Staining with supernatants from untransfected cells (no plasmid) is used as a negative control. All data are normalized to mode. **(B)** Table with the antibodies presented in panel a, showing the heavy and light chain V genes. The (-) symbol indicates no binding to the cognate cell line. The intermediate blue lines separate the data according to cell line (VSG3$_{WT}$, VSG3$_{S319A}$, VSG3$_{S317A}$, VSG3$_{SSAA}$). The CDR3s from each antibody is shown along with the original antibody isotype and somatic hypermutations, if any. **(C)** FACS histograms showing the binding to S317A-covered trypanosomes of different recombinant antibodies, sharing the same gn33 (same V, J and CDR3) light chain but replacing the heavy chain with V*H* genes that are present in the repertoires but not as pairs with Vκ gn33. The blue color indicates binding and the red non-binding. Staining with supernatant from untransfected cells is used as a negative control (black line). All FACS data shown are normalized to mode. **(D)** Table showing the same data as in panel a. V and J segments as well as the CDR3 of both heavy and light are shown, along with the binding to the individual cell lines and the original isotypes. The (+) symbol indicates binding while the (-) lack of binding. The original antibody isotype, as well as somatic hypermutations, if any, are also shown.

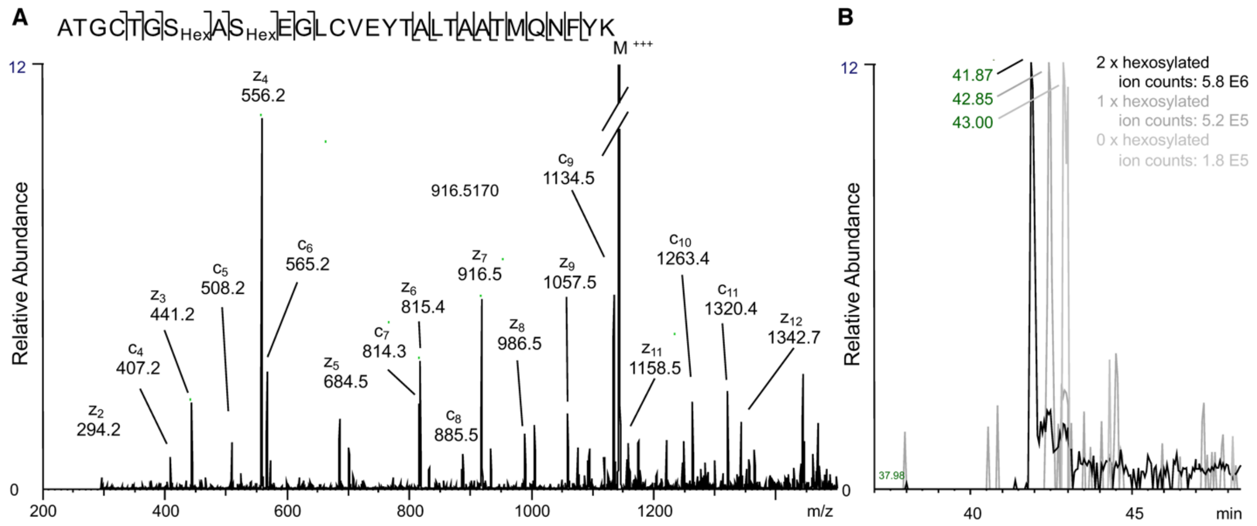

**Figure S7: The VSG3 peptide A311-K339 carrying two hexose residues is the dominant species compared to those carrying fewer hexoses. (A)** The triply charged precursor mass of the VSG3 peptide A311-K339 modified by two hexose residues is fragmented by electron transfer dissociation, which preserves the position of posttranslational modification. S317 and S319 are both mono hexosylated. **(B)** An extracted ion chromatogram (XIC) showing the elution profiles of the triply charged VSG3 peptides without hexosylation (m/z=1040.132) and modified by one (m/z= 1094.149) or two (m/z=1148.172) hexoses. Peak height is set to 100 for each XIC for better visibility. The absolute ion count for each peptide is indicated.

| | $VSG2_{WT}$ | $VSG2_{AAA}$ |
|---|---|---|
| **Data Collection** | | |
| Beamline | APS 24-ID-C | SLS X06DA (PXIII) |
| Processing software | RAPD | go.pi |
| Wavelength (Å) | 0.9791 | 1.0 |
| Resolution range (Å) | 61.81 -1.735 (1.797 -1.735) | 48.01 -1.961 (2.031-1.961) |
| Space group | P 1 21 1 | P 41 21 2 |
| Unit cell a, b, c (Å) | 42.915 87.585 88.121 | 96.015 96.015 108.418 |
| Unit cell α, β, γ (°) | 90 98.135 90 | 90 90 90 |
| Total reflections | 218133 (19957) | 721544 (12060) |
| Unique reflections | 65518 (5809) | 36675 (3341) |
| Multiplicity | 3.3 (3.2) | 19.7 (3.6) |
| Completeness (%) | 97.26 (86.82) | 99.24 (92.39) |
| Mean I/sigma(I) | 10.93 (0.75) | 20.27 (0.84) |
| Wilson B-factor | 27.53 | 34.20 |
| *R-merge* | 0.07673 (1.565) | 0.09816 (0.8934) |
| *R-meas* | 0.09159 (1.882) | 0.1006 (1.023) |
| *R-pim* | 0.04939 (1.032) | 0.02123 (0.4747) |
| CC1/2 | 0.998 (0.307) | 0.999 (0.557) |
| CC* | 0.999 (0.685) | 1 (0.846) |
| **Refinement** | | |
| Refinement reflections | 64942 (5809) | 36672 (3341) |
| R-free reflections | 3206 (289) | 1837 (167) |
| R-work | 0.1887 (0.3309) | 0.1637 (0.3229) |
| R-free | 0.2209 (0.3402) | 0.1970 (0.3435) |
| CC(work) | 0.961 (0.650) | 0.972 (0.716) |
| CC(free) | 0.943 (0.611) | 0.899 (0.748) |
| Number of non-hydrogen atoms | 5360 | 2953 |
| macromolecules | 4928 | 2664 |
| ligands | 91 | 50 |
| solvent | 341 | 239 |
| Protein residues | 686 | 363 |
| RMS(bonds) | 0.006 | 0.009 |
| RMS(angles) | 0.76 | 1.12 |
| Ramachandran favored (%) | 97.01 | 97.20 |
| Ramachandran allowed (%) | 2.99 | 2.80 |
| Ramachandran outliers (%) | 0.00 | 0.00 |
| Rotamer outliers (%) | 0.00 | 0.00 |
| Clashscore | 0.91 | 1.11 |
| Average B-factor | 34.79 | 43.90 |
| macromolecules | 34.24 | 43.44 |
| ligands | 46.47 | 58.31 |
| solvent | 39.65 | 46.03 |
| Number of TLS groups | 40 | 10 |

Highest-resolution shell statistics are in parentheses.

**Table S1: Crystallographic statistics for $VSG2_{WT}$ and AAA-mutant.**

| | VSG3$_{WT}$ | VSG3$_{S317A}$ | VSG3$_{S319A}$ | VSG3$_{SSAA}$ |
|---|---|---|---|---|
| **Data Collection** | | | | |
| Beamline | SLS X06DA (PXIII) | SLS X06DA (PXIII) | Diamond i03 | SLS X06DA (PXIII) |
| Processing software | go.pi | go.pi | Xia2 Dials | go.pi |
| Wavelength (Å) | 1.0 | 1.0 | 0.9763 | 1.0 |
| Resolution range (Å) | 40.84-1.273 (1.318-1.273) | 45.75 - 1.95 (2.02 - 1.95) | 40.8 - 1.13 (1.17 - 1.13) | 40.85 - 1.423 (1.474 - 1.423) |
| Space group | I 21 3 | I 21 3 | I 21 3 | I 21 3 |
| Unit cell a, b, c (Å) | 129.155 129.155 129.155 | 129.396 129.396 129.396 | 129.007 129.007 129.007 | 129.183 129.183 129.183 |
| Unit cell α, β, γ (°) | 90 90 90 | 90 90 90 | 90 90 90 | 90 90 90 |
| Total reflections | 1858461 (178155) | 812357 (13521) | 5167710 (373961) | 1340137 (127475) |
| Unique reflections | 93285 (9281) | 26348 (2584) | 132433 (8709) | 66947 (6654) |
| Multiplicity | 19.9 (19.1) | 30.8 (5.2) | 39.0 (28.4) | 20.0 (19.1) |
| Completeness (%) | 99.94 (99.66) | 99.89 (99.12) | 96.60 (66.13) | 99.95 (99.64) |
| Mean I/sigma(I) | 21.27 (1.28) | 32.36 (1.67) | 23.97 (0.42) | 20.08 (1.19) |
| Wilson B-factor | 17.02 | 24.43 | 17.40 | 20.10 |
| *R-merge* | 0.08937 (2.497) | 0.1179 (0.8387) | 0.07786 (6.968) | 0.1164 (2.767) |
| *R-meas* | 0.0917 (2.565) | 0.1198 (0.9312) | 0.07886 (7.094) | 0.1195 (2.843) |
| *R-pim* | 0.02045 (0.5849) | 0.0209 (0.3875) | 0.0125 (1.324) | 0.02664 (0.6492) |
| CC1/2 | 1 (0.513) | 0.999 (0.614) | 1 (0.169) | 1 (0.455) |
| CC* | 1 (0.823) | 1 (0.872) | 1 (0.538) | 1 (0.791) |
| **Refinement** | | | | |
| Refinement reflections | 93232 (9276) | 26341 (2581) | 127939 (8709) | 66920 (6653) |
| R-free reflections | 4662 (464) | 1318 (129) | 6420 (411) | 3346 (333) |
| R-work | 0.1775 (0.3804) | 0.1816 (0.2947) | 0.1755 (0.3403) | 0.1742 (0.2958) |
| R-free | 0.1983 (0.3865) | 0.2190 (0.3349) | 0.1981 (0.3482) | 0.1928 (0.3273) |
| CC(work) | 0.479 (0.036) | 0.955 (0.785) | 0.966 (0.512) | 0.961 (0.727) |
| CC(free) | 0.482 (-0.004) | 0.927 (0.666) | 0.963 (0.480) | 0.953 (0.693) |
| Number of non-hydrogen atoms | 3013 | 2838 | 3177 | 2942 |
| macromolecules | 2622 | 2529 | 2597 | 2624 |
| ligands | 94 | 72 | 83 | 72 |
| solvent | 297 | 237 | 497 | 246 |
| Protein residues | 361 | 361 | 363 | 366 |
| RMS(bonds) | 0.017 | 0.010 | 0.018 | 0.006 |
| RMS(angles) | 1.51 | 0.94 | 1.44 | 0.97 |
| Ramachandran favored (%) | 98.03 | 97.16 | 97.73 | 97.46 |
| Ramachandran allowed (%) | 1.97 | 2.84 | 2.27 | 2.26 |
| Ramachandran outliers (%) | 0.00 | 0.00 | 0.00 | 0.28 |
| Rotamer outliers (%) | 0.00 | 0.00 | 0.00 | 0.00 |
| Clashscore | 0.19 | 0.00 | 0.57 | 0.57 |
| Average B-factor | 22.25 | 24.60 | 24.11 | 24.97 |
| macromolecules | 21.45 | 23.97 | 22.30 | 24.28 |
| ligands | 24.21 | 34.39 | 29.19 | 28.91 |
| solvent | 28.76 | 28.39 | 32.75 | 31.18 |
| Number of TLS groups | 6 | 4 | 6 | 6 |

Highest-resolution shell statistics are in parentheses.

**Table S2: Crystallographic statistics for VSG3$_{WT}$ and the sugar-mutants.**

| Protein Sample | # Variable Gal† | # $Ca^{2+}$ bound | Mass (Da) | | |
|---|---|---|---|---|---|
| | | | Predicted‡ | Measured§ | Δ |
| **$VSG2_{WT}$, No EDTA** | 1 | 0 | 95.371,5 | 95.373,6 | 2,1 |
| | | 1 | 95.409,6 | 95.409,2 | -0,4 |
| | | 2 | 95.447,7 | 95.446,2 | -1,5 |
| | 2 | 0 | 95.533,7 | 95.532,2 | -1,5 |
| | | 1 | 95.571,7 | 95.570,2 | -1,5 |
| | | 2 | 95.609,8 | 95.610,0 | 0,2 |
| | 3 | 0 | 95.695,8 | 95.694,0 | -1,8 |
| | | 1 | 95.733,9 | 95.732,4 | -1,5 |
| | | 2 | 95.771,9 | 95.771,4 | -0,5 |
| | 4 | 0 | 95.858,0 | 95.856,4 | -1,6 |
| | | 1 | 95.896,0 | 95.894,4 | -1,6 |
| | | 2 | 95.934,1 | 95.933,4 | -0,7 |
| | 5 | 0 | 96.020,1 | 96.018,0 | -2,1 |
| | | 1 | 96.058,2 | 96.056,8 | -1,4 |
| | | 2 | 96.096,2 | 96.095,4 | -0,8 |
| | 6 | 0 | 96.182,2 | 96.180,8 | -1,4 |
| | | 1 | 96.220,3 | 96.219,2 | -1,1 |
| | | 2 | 96.258,4 | 96.257,8 | -0,6 |
| | 7 | 0 | 96.344,4 | 96.343,2 | -1,2 |
| | | 1 | 96.382,4 | 96.380,6 | -1,8 |
| | | 2 | 96.420,5 | 96.419,4 | -1,1 |
| | 8 | 0 | 96.506,5 | 96.503,4 | -3,1 |
| | | 1 | 96.544,6 | 96.543,6 | -1,0 |
| | | 2 | 96.582,7 | 96.583,2 | 0,5 |
| **$VSG2_{WT}$, +EDTA** | 1 | 0 | 95.371,5 | 95.370,8 | -0,7 |
| | 2 | 0 | 95.533,7 | 95.531,8 | -1,9 |
| | 3 | 0 | 95.695,8 | 95.694,0 | -1,8 |
| | 4 | 0 | 95.858,0 | 95.855,4 | -2,6 |
| | 5 | 0 | 96.020,1 | 96.017,6 | -2,5 |
| | 6 | 0 | 96.182,2 | 96.180,2 | -2,0 |
| | 7 | 0 | 96.344,4 | 96.343,2 | -1,2 |
| | 8 | 0 | 96.506,5 | 96.505,6 | -0,9 |
| | 9 | 0 | 96.668,7 | 96.670,2 | 1,5 |
| **$VSG2_{AAA}$, No EDTA** | 0 | 0 | 94.947,3 | 94.945,6 | -1,7 |
| | 1 | 0 | 95.109,4 | 95.106,6 | -2,8 |
| | 2 | 0 | 95.271,6 | 95.269,6 | -2,0 |
| | 3 | 0 | 95.433,7 | 95.431,4 | -2,3 |
| | 4 | 0 | 95.595,9 | 95.593,8 | -2,1 |
| | 5 | 0 | 95.758,0 | 95.756,0 | -2,0 |
| | 6 | 0 | 95.920,1 | 95.918,2 | -1,9 |
| | 7 | 0 | 96.082,3 | 96.079,0 | -3,3 |
| | 8 | 0 | 96.244,4 | 96.242,4 | -2,0 |
| | 9 | 0 | 96.406,6 | 96.404,4 | -2,2 |

* Only peak assignments for the full N-deglycosylated VSG2 dimers are shown.
† After complete PNGaseF N-deglycosylation, the remaining glycan microheterogeneity in VSG2 comes from variable galactosylation at the C-terminal GPI anchor.
‡ See the Methods section for details on how the predicted masses were calculated.
§ From deconvolution of nMS spectra using the UniDec software.

**Table S3: Native MS analyses of the PNGaseF-treated VSG2 samples.***
